# Concurrent AXL Inhibition Enhances RAS and ERK Inhibitor Efficacy in *KRAS*-mutant Pancreatic and Lung Cancer

**DOI:** 10.64898/2026.08.10.743026

**Authors:** Yan Ming Ching, Shruthi Narayanan, Jeffrey A. Klomp, Tamara Isermann, Sina Loewe, Wen-Hsuan Chang, Andrew M. Waters, Sheila R. Nicewarner Peña, Elisa Baldelli, A. Cole Edwards, Teresa Börding, Runying Yang, Craig M. Goodwin, Prson Gautam, Mariano Ponz-Sarvisé, David Horst, Kyle Seamon, Yongxian Zhuang, Linh Tran, Jingjing Jiang, Mallika Singh, Krister Wennerberg, Emanuel F. Petricoin, Kirsten L. Bryant, Clint A. Stalnecker, H. Shelton Earp, Adrienne D. Cox, Christine Sers, Silvestre Vicent, Channing J. Der, Bjoern Papke

**Affiliations:** Institute of Pathology, Charité – University Medical Center Berlin, corporate member of Freie Universität Berlin and Humboldt-Universität zu Berlin; Center for Applied Medical Research, Program in Solid Tumors, University of Navarra, Pamplona, Spain; Medical Oncology Department, Cancer Center Clínica Universidad de Navarra, Pamplona, Spain; Department of Medicine, College of Human Medicine, Michigan State University, Grand Rapids, Michigan; German Cancer Consortium (DKTK) Partner Site Berlin, German Cancer Research Center (DKFZ), Heidelberg, Germany; Lineberger Comprehensive Cancer Center, University of North Carolina at Chapel Hill, Chapel Hill, North Carolina; Department of Surgery, University of Cincinnati College of Medicine, Cincinnati, Ohio; Department of Cancer Biology, University of Cincinnati, Cincinnati, Ohio; Center for Applied Proteomics and Molecular Medicine, George Mason University, Manassas, Virginia; Institute for Molecular Medicine Finland, University of Helsinki, Helsinki, Finland; Navarra Institute for Health Research (IdiSNA), Pamplona, Spain; Revolution Medicines, Redwood City, California; Biotech Research and Innovation Centre (BRIC) and Novo Nordisk Foundation Center for Stem Cell Biology (DanStem), University of Copenhagen, Copenhagen, Denmark; Department of Pharmacology, University of North Carolina at Chapel Hill, Chapel Hill, North Carolina; Department of Biochemistry and Biophysics, University of North Carolina at Chapel Hill, Chapel Hill, North Carolina; Department of Medicine, Division of Oncology, University of North Carolina at Chapel Hill, Chapel Hill, North Carolina; Department of Radiation Oncology, University of North Carolina at Chapel Hill, Chapel Hill, North Carolina; Centro de Investigación Biomédica en Red de Cáncer (CIBERONC), Madrid, Spain; Department of Pathology, Anatomy, and Physiology, University of Navarra, Pamplona, Spain

**Author notes:** **Corresponding Authors:** Christine Sers, Institute of Pathology, Charité - Universitätsmedizin Berlin, Berlin, Germany.; Silvestre Vicent, 55 Pio XII Ave., 31008, Pamplona, Spain. Channing J. Der, Lineberger Comprehensive Cancer Center, University of North Carolina at Chapel Hill, Chapel Hill, NC 27599. Bjoern Papke, Institute of Pathology, Charité - Universitätsmedizin Berlin, Berlin, Germany. Y.M. Ching and S. Narayanan are co-first authors of this study. C. Sers, S. Vicent, C.J. Der, and B. Papke are co-senior authors of this article.

**Keywords:** KRAS, RAS, ERK, AXL, RAS(ON) multi-selective inhibitor, PDAC

## Abstract

Resistance limits the clinical efficacy of RAS inhibitors. We applied chemical and genetic screens and identified the AXL receptor tyrosine kinase as a driver of resistance to RAS-ERK inhibition. We determined that combination treatment with the AXL inhibitor bemcentinib (AXLi) together with the RAS(ON) multi-selective tri-complex inhibitor RMC-7977 (RASi) or the ERK-selective inhibitor SCH772984 (ERKi) significantly enhanced growth suppression in human *KRAS*-mutant pancreatic and lung cancer models. Combined AXLi and RASi treatment of human *KRAS*-mutant pancreatic cell line−derived xenograft tumors synergistically suppressed ERK activation and MYC expression, and caused tumor regression. Analyses of immunocompetent mouse allograft pancreatic tumor models revealed a largely tumor cell−intrinsic response to inhibitor treatment. We identified an unexpected mechanism whereby KRAS inhibition upregulated the AXL ligand GAS6, activating AXL but inducing an AXL-dependent adaptive resistance mechanism wherein AXL antagonizes RASi efficacy. Our observations support concurrent AXL inhibition as a strategy to enhance RAS inhibitor clinical efficacy.

**STATEMENT OF SIGNIFICANCE:** Our findings identify AXL as a driver of resistance to RAS inhibitors, establishing a combination strategy to overcome resistance and enhance RAS inhibitor therapeutic efficacy in *KRAS*-mutant cancer by maximally inhibiting oncogenic RAS signaling.

## INTRODUCTION

Pancreatic cancer is the third leading cause of cancer-related deaths in the United States, with a dismal 5-year survival rate of approximately 13% (1). Despite the well-defined genetic landscape of pancreatic ductal adenocarcinoma (PDAC) (2), the predominant form of pancreatic cancer (3), no effective targeted therapies have been developed, and current standards-of-care are ineffective cytotoxic drugs (4). A defining feature of PDAC is the near-universal occurrence of and dependency on oncogenic *KRAS* mutations (5), which are present in approximately 95% of cases (6). *KRAS* mutations are both the initiating genetic event and essential for tumor maintenance, thereby establishing KRAS as a critical therapeutic target in PDAC (5).

Of the remaining 5% of PDAC that are wild-type for KRAS, 60% harbor mutant genes encoding components that cause activation of the key KRAS effector pathway (6), the RAF-MEK-ERK mitogen-activated protein kinase (MAPK) cascade. This observation, together with studies demonstrating that aberrant ERK activation alone can phenocopy mutant KRAS-driven PDAC development and growth (7–10), supports the critical role of ERK MAPK signaling in driving PDAC growth.

Recent advancements in directly targeting mutant KRAS have led to the approval of the first KRAS inhibitors for the treatment of non-small cell lung and colorectal cancers (11–13). This success has fueled the development and clinical evaluation of additional RAS inhibitors that target virtually every KRAS mutation. In particular, we showed that the RAS(ON) multi-selective tri-complex inhibitor tool compound RMC-7977 (8,14) and the related clinical candidate analog daraxonrasib/RMC-6236 (15) exhibited potent anti-tumor activity against a spectrum of non-KRAS^G12C^-mutant PDAC and other cancers. Recently we reported phase 3 clinical trial results in PDAC of second-line treatment with daraxonrasib, where overall survival (OS) was nearly double that seen with standard-of-care chemotherapy (16), supporting ongoing clinical evaluation as first line treatment for PDAC (NCT07491445).

However, even with this favorable clinical activity, the limited overall response rate and OS reflect the same challenges of primary/innate and acquired resistance encountered previously with KRAS^G12C^ inhibitors (17,18). These findings underscore the widely accepted belief that combination strategies will be pivotal for the development of more durable RAS inhibitor-based therapies for *KRAS*-mutant cancers. Reflecting this belief, ongoing clinical trials are evaluating various RAS inhibitors in combination with inhibitors of other RAS signaling components (e.g., SHP2, MEK), immune checkpoint inhibitors and chemotherapy (11–13).

The predominant mechanism of acquired resistance to RAS inhibition is ERK reactivation (12,19). In this study, we aimed to identify rational combination strategies that enhance the efficacy of RAS-ERK MAPK pathway inhibition in *KRAS*-mutant PDAC. We applied two independent functional screening strategies that together identified AXL, a member of the TAM family (TYRO3, AXL, MERTK) of receptor tyrosine kinases (RTKs) (20–22) as a driver of resistance to RAS and ERK inhibitor treatment. We observed that inhibition of RAS or ERK triggered a compensatory feedback response characterized by increased expression of the AXL ligand GAS6. Moreover, concurrent treatment with the RAS(ON) multi-selective tri-complex inhibitor RMC-7977 and the AXL-selective inhibitor bemcentinib (BGB324) caused synergistic growth suppression of PDAC and lung adenocarcinoma (LUAD) *in vitro* and induced regression of tumors *in vivo*. Unexpectedly, given the significant role that AXL serves in immune cell function (23–25), this combination showed the same synergistic anti-tumor activity in a syngeneic immunocompetent mouse model of *Kras*^G12D^-mutant PDAC. Consistent with this, our evaluation of immune cell function in tumors from treated mice indicated a largely tumor cell−intrinsic response to the combination. Finally, we identified an unexpected mechanism for AXL-driven RAS inhibitor resistance that was independent of conventional AXL effector signaling. Instead, we found that RAS inhibition induced compensatory AXL activation, which then attenuated RAS inhibitor efficacy in suppressing ERK and MYC activation. In summary, our findings support the potential of combining AXL inhibition with RAS-targeted therapies to enhance antitumor efficacy and improve clinical outcomes in *KRAS*-mutant PDAC and LUAD.

## RESULTS

### AXL Suppression Enhances Sensitivity to KRAS and ERK Inhibition in PDAC

To identify combination strategies to improve therapeutic targeting of aberrant KRAS activation and ERK MAPK signaling in PDAC, we first performed drug sensitivity and resistance testing (DSRT) on a panel of 20 *KRAS*-mutant PDAC cell lines. We applied a chemical library comprised of 525 U.S. Food and Drug Administration/European Medicines Agency (FDA/EMA)-approved or clinical candidate oncology drugs (26) to identify combinations that enhanced the growth inhibitory activity of the ERK-selective inhibitor SCH772984 (ERKi) (27) (Supplementary Fig. S1A).

Consistent with our previous observations (28–31), we identified inhibitors of signaling components within the RAS signaling network that further enhanced ERKi growth suppression. These included vertical pathway inhibition of EGFR, RAF and MEK, inhibitors of PI3K-AKT-mTOR signaling, and SRC and IGFR1 inhibitors (Fig. 1A). Additional drugs with enhanced activities included inhibitors of HDAC (quisinostat and rocilinostat) and Aurora kinase (PF-03814735, ENMD-2076 and danusertib), the multi-RTK inhibitor nintedanib, the PAK inhibitor FRAX486 and the AXL inhibitor (AXLi) bemcentinib/BGB324. The responses to all inhibitors were heterogeneous, in that a subset of cell lines was responsive to each combination. For example, cotreatment with AXLi further reduced proliferation in response to ERKi in 13 of the 20 PDAC cell lines (Fig. 1B).

**Figure 1.**
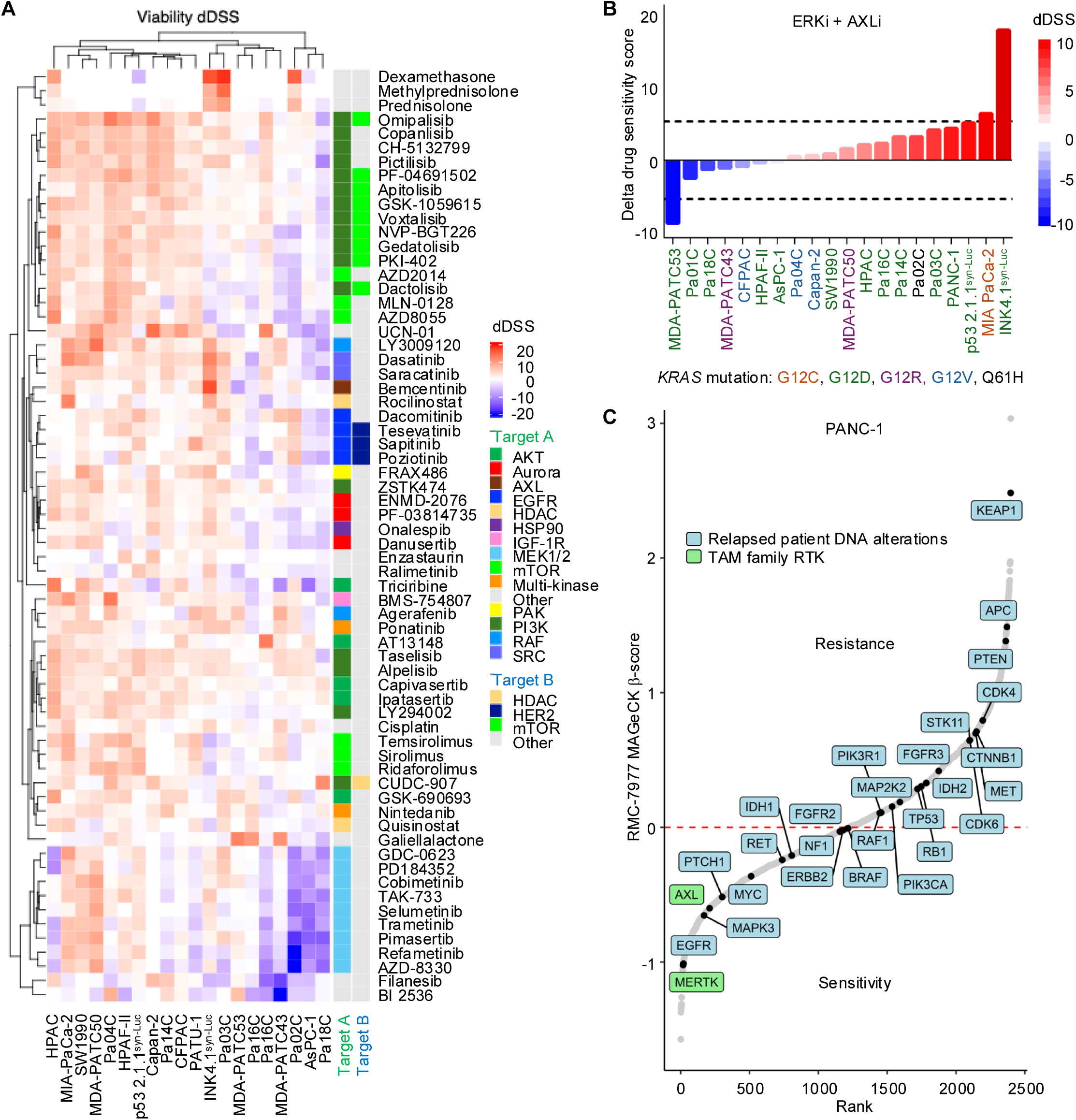
DSRT chemical library and CRISPR genetic loss-of-function screens converge on AXL loss as a sensitizer to RAS-ERK MAPK inhibition. **A,** Clustered heatmap of delta drug sensitivity scores (dDSS) from the DSRT viability screen across (columns) a panel of 18 human and two mouse (*Kras*^G12D^; *Ink4a*^null^ or *Tp53*^null^) (7) PDAC cell lines, in the presence of the ERK1/2-selective inhibitor SCH772984. Each row represents an individual drug. Color indicates the dDSS (red, increased synergy; blue, decreased; white, no change). Rows and columns are hierarchically clustered (dendrograms). **B,** Waterfall plot illustrating combination viability scores from a DSRT screen using a 525-oncology drug library in the presence of 100 nM SCH772984 (ERKi). The plot displays the dDSS values for bemcentinib (AXLi) in combination with ERKi for a panel of PDAC cell lines. Red indicates additive (between 0-5) and synergistic (> 5) effects, blue indicates antagonistic (< additive) effects, and white indicating no significant change. Cell line color labels indicate the KRAS mutation. **C,** Snake plot of genes ranked by enrichment or depletion (delta beta scores) from a CRISPR-Cas9 loss-of-function screen in PANC-1 cells following four weeks of treatment with the RAS(ON) multi-selective inhibitor RMC-7977 (RASi), relative to DMSO control. Highlighted genes correspond to relapsed patient DNA alterations (blue) or TAM family RTK (green).

In our clinical evaluation of ERK inhibitors in PDAC, we found that efficacy was hampered by significant on-target toxicity (32,33). By comparison, although we demonstrated recently that KRAS drives PDAC gene transcription and growth predominantly through ERK (9,10), KRAS^G12C^ inhibitors have shown both strong efficacy and acceptable toxicity (11–13). Therefore, we next evaluated combinations with the RAS(ON) multi-selective tri-complex inhibitor RMC-7977 (designated RASi) (8,14).

To identify genetic modulators of responses to RASi, we performed a CRISPR-Cas9 loss-of-function screen in the *KRAS*^G12D^-mutant PDAC cell line PANC-1, using a druggable genome library targeting 2,240 genes that encode components of major cancer signaling pathways (29,34) (Supplementary Fig. S1B). Supporting the ability of this screen to identify *bona fide* genetic regulators of RAS inhibitor sensitivity and resistance, we detected 26 genes previously identified as altered in patients who relapsed on KRAS^G12C^ inhibitor treatment (35–39) (Fig. 1C). For example, sgRNAs targeting tumor suppressor genes (e.g., *KEAP1*, *APC*, *PTEN*, and *RB1*) identified in such patients were among the sgRNAs enriched in our screen and sgRNAs targeting genes found to be mutationally activated (e.g., *EGFR*) or amplified (e.g., *MYC*) in relapsed patients (35–37,39) were among the depleted sgRNAs. Notably, our screen additionally demonstrated that genetic knockout of *AXL* sensitized PANC-1 cells to RASi treatment (Fig. 1C). This finding is consistent both with our chemical library screen (Fig. 1A) and with our previous CRISPR screen using a smaller 384-oncology gene library, where we detected that loss of *AXL* increased sensitivity to MEK and ERK inhibitors in a panel of *KRAS*-mutant PDAC, and to a lesser degree in lung and colorectal carcinoma cell lines (40). Collectively, these findings from distinct independent unbiased screens strongly support AXL as a promising co-target to enhance the therapeutic efficacy of RAS-ERK inhibitors in PDAC.

### AXL Inhibition Suppresses PDAC Growth

We next assessed the potential role of AXL as a cancer driver in PDAC. First, we investigated the clinical relevance of *AXL* mRNA expression in PDAC tumors by examining patient survival data from The Cancer Genome Atlas (TCGA) dataset (Fig. 2A). Our findings indicate that tumors exhibiting high *AXL* expression are significantly correlated with reduced overall survival.

**Figure 2.**
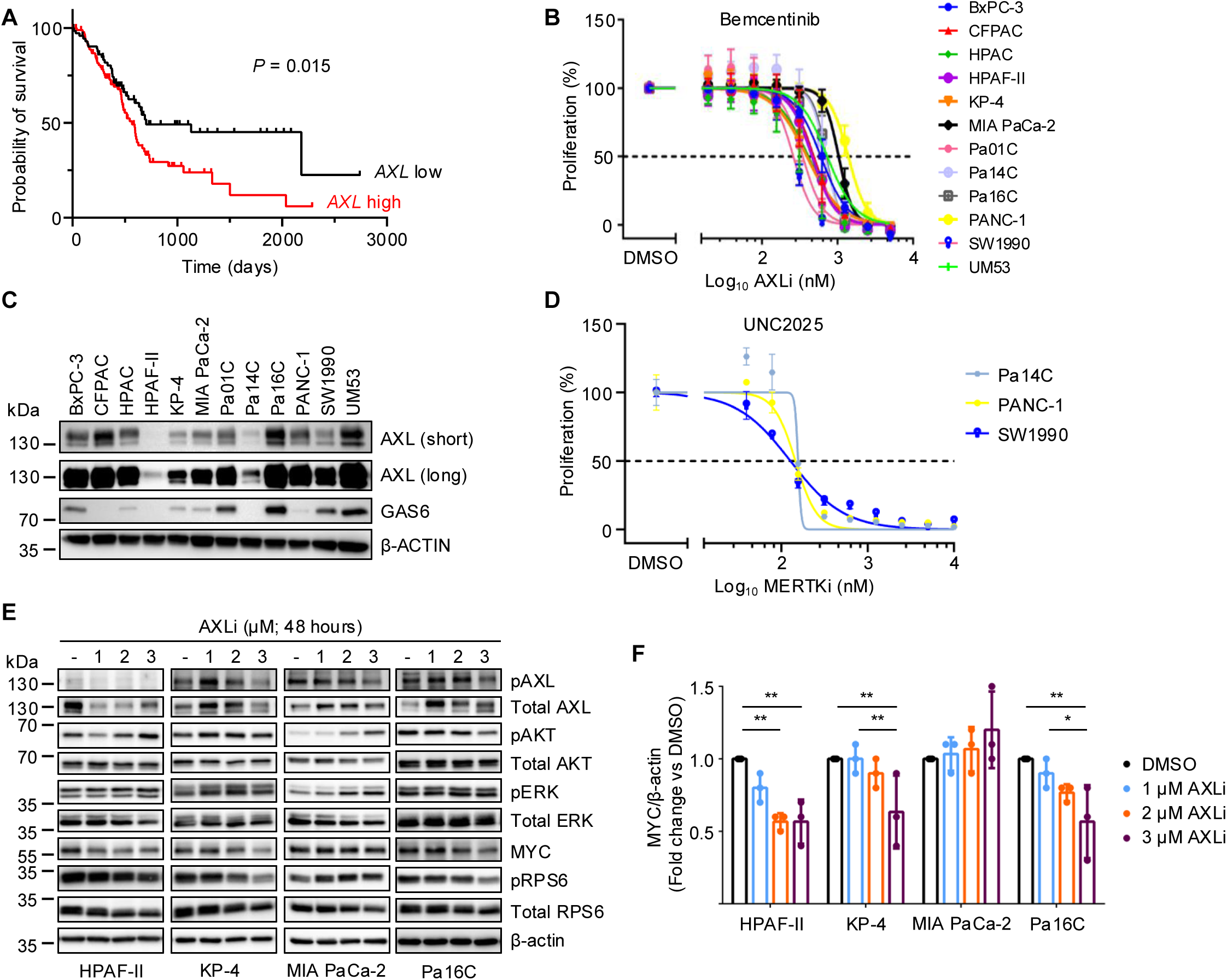
AXL inhibition suppresses PDAC growth *in vitro*. **A,** Kaplan-Meier survival curves of PDAC patients stratified by high versus low *AXL* expression (data from the TCGA database). Statistical significance between high versus low *AXL* expression was determined by log-rank test (*P* = 0.015). **B,** Western blot analysis of AXL (short and long exposure time) and GAS6 expression in a panel of PDAC cell lines (*n* = 3 biological replicates). b-actin expression was determined to verify equivalent total protein. **C,** Proliferation of PDAC cell lines treated with increasing concentrations of AXLi for 5 days, measured as percent proliferation relative to DMSO control. The dashed line indicates 50% growth inhibition (GI_50_). Data represent three independent biological replicates (*n* = 3) ± SD. **D,** Proliferation of Pa14C, PANC-1, and SW1990 cells treated with increasing concentrations of the MERTK inhibitor UNC2025 for 5 days, measured as percent proliferation relative to DMSO control. The dashed line indicates GI_50_. Points are the mean ± SD (*n* = 3 biological replicates). **E,** Western blot analysis of signaling components downstream of AXL to monitor target inhibition in a panel of PDAC cells treated with increasing doses of AXLi (1, 2 and 3 µM) for 48 hours. (*n* = 3 biological replicates). **F,** Quantification of MYC expression from the Western blots shown in panel E, normalized to β-actin (loading control) and expressed relative to DMSO. Data shown as mean ± SD. Statistical significance between treatments and DMSO control was determined by one-way ANOVA (\**P <* 0.05, \*\**P <* 0.01). Only statistically significant results are shown.

Since AXL has a well-established role as an oncogene (24,25,41), we next set out to investigate the therapeutic potential of targeting AXL in PDAC. We explored the *in vitro* sensitivity of a panel of 12 PDAC cell lines to the clinical candidate AXL-selective inhibitor bemcentinib/BGB324 (42) as monotherapy. We observed variable sensitivity, with GI₅₀ values ranging from 260 to 1386 nM (Fig. 2B; Supplementary Fig. S2A and S2B). This growth inhibitory activity is comparable with that seen in previous analyses of human and mouse PDAC cell lines (43). Unexpectedly, AXLi sensitivity did not correlate with baseline levels of AXL or its ligand GAS6 (Supplementary Fig. S2B), nor did we detect a significant correlation between receptor and ligand at either the protein or mRNA level (Fig. 2C; Supplementary Fig. S2C and S2D).

From our CRISPR screen, we also identified a second TAM family RTK, *MERTK* (Fig. 1C), which shares significant functional relationships with AXL (20–22). We therefore assessed the role of MERTK in PDAC growth. We treated PDAC cell lines with UNC2025, a potent MERTK-selective inhibitor with 12- to 15-fold lower activity against AXL and TYRO3, respectively (44). In contrast, bemcentinib is AXL-selective, with greater than 100-fold selectivity over MERTK and TYRO3 as well as other RTKs (42). However, UNC2025 affinity for AXL is equivalent to that of bemcentinib. We observed that UNC2025 was significantly more potent than bemcentinib, exhibiting 120 −150 nM GI_50_ activities (Fig. 2D; Supplementary Fig. S2E), presumably due to the potent inhibition of both MERTK and AXL.

We next determined if the degree of AXLi sensitivity was associated with the degree of target inhibition. For these analyses, we evaluated the phosphorylation of components of AXL signaling shown previously to be blocked by AXLi in PDAC cell lines *in vitro* (43) (Supplementary Fig. S2F). Following treatment of PDAC cells for 48 hours with increasing doses of AXLi, we found variable and inconsistent reductions in AXL autophosphorylation at residue Y779, a direct marker of AXL kinase activation. We also evaluated downstream effector signaling and phosphorylation of ERK (T202/Y204), AKT (S473) and ribosomal protein S6 (S235/236; pRPS6). Surprisingly, although there were evident cell line–specific differences, we observed no significant and consistent reduction across pERK or pAKT (Fig. 2E). pRPS6 levels did show a modest dose-dependent decrease in three of four cell lines. Finally, previous studies implicated AXL signaling in the upregulation of MYC expression (45,46). We also observed a dose-dependent downregulation of MYC in three of four cell lines, somewhat more robustly than in pRPS6, thereby providing perhaps the most consistent marker associated with AXLi treatment (Fig. 2E and 2F).

### Concurrent Inhibition of AXL and RAS-ERK Synergistically Impairs PDAC Proliferation *In Vitro*

To further investigate the molecular response of PDAC cell lines to RAS-ERK pathway inhibition, we assessed the levels of AXL and its ligand GAS6 following treatment with either RASi or ERKi for 48 hours. We observed a consistent upregulation of GAS6 across four PDAC cell lines in response to RAS-ERK inhibition, suggesting compensatory feedback activation of AXL (Fig. 3A). To further validate these findings, we retrospectively analyzed our recently described KRAS-ERK gene signature in PDAC (8). Expression of *DUSP6*, a well-validated ERK-regulated gene, was reduced upon genetic depletion of *KRAS* and upon pharmacological inhibition of mutant KRAS or ERK (Fig. 3B). In contrast, both approaches increased *GAS6* expression whereas *AXL* remained unchanged. These observations suggest that KRAS-ERK inhibition–induced GAS6 upregulation may be one mechanistic basis for our identification of *AXL* as a gene that modulates sensitivity to RAS inhibition.

**Figure 3.**
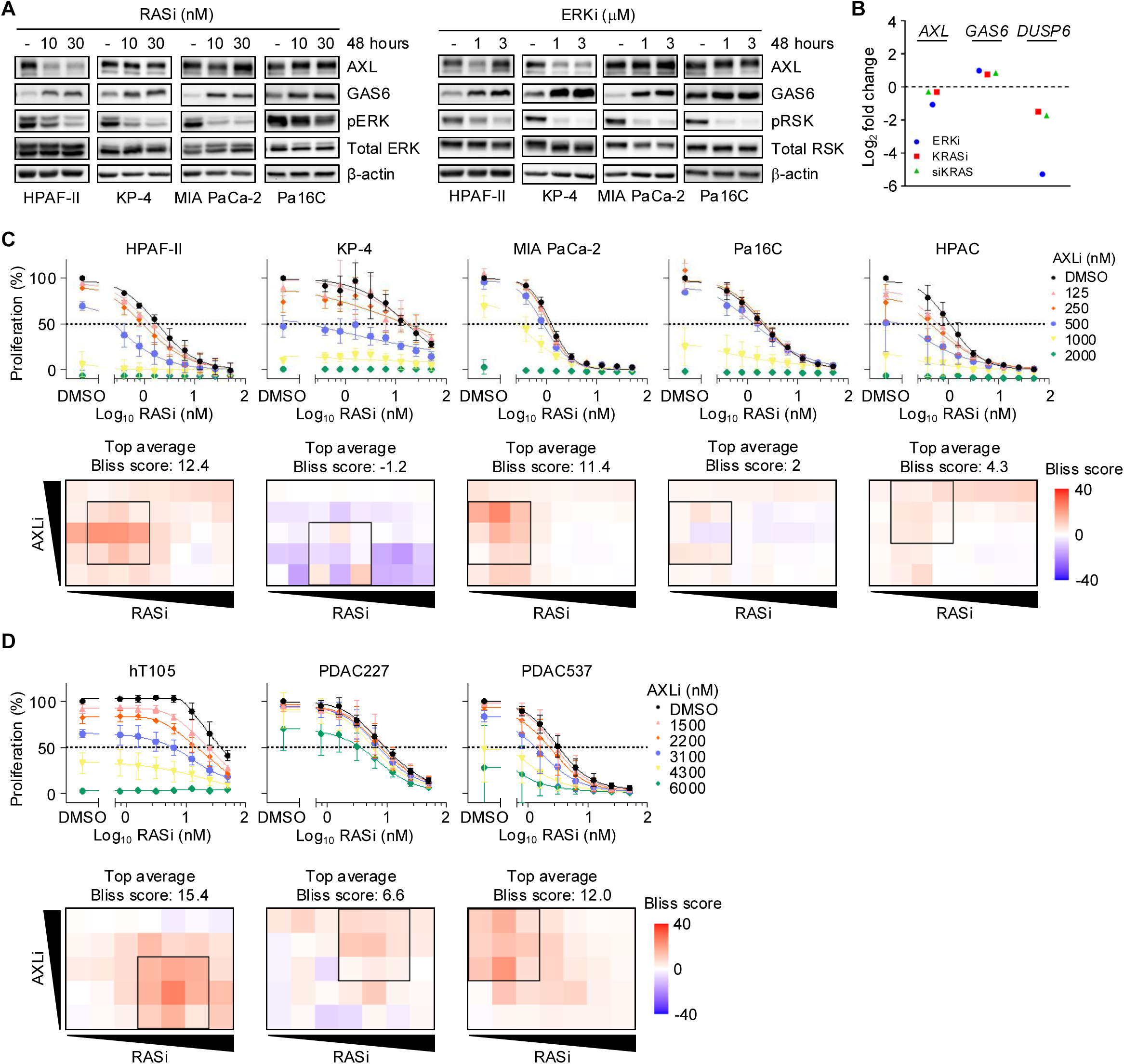
RASi treatment causes GAS6 upregulation and when combined with AXLi synergistically inhibits PDAC growth. **A,** Western blot evaluation of AXL, GAS6 and pERK expression in PDAC cells treated with DMSO and RAS inhibitor (10 and 30 nM) (left panel) or ERK inhibitor (1 and 3 µM) (right panel) for 48 hours (*n* = 3 biological replicates). **B**, RNA-sequencing analysis highlighting transcriptional changes in *GAS6* and *AXL* following ERK inhibition (blue circles), KRAS knockdown (green triangles) (both in PDAC cell lines), or KRAS inhibition (red squares) after 24 hours in three KRAS^G12C/D^-mutant human PDAC cell line-derived tumor xenografts (CDX) models. *DUSP6* was included as a control for RAS-ERK target inhibition (data reported previously in (9)). **C,** Upper panels: Proliferation of PDAC cell lines treated with increasing concentrations of RASi and AXLi (125-2000 nM) for 5 days, shown as percent proliferation relative to DMSO; the dashed line indicates GI_50_ (*n* = 3 biological replicates). Lower panels: BLISS synergy heatmaps (red, synergy; blue, antagonism; scale −40 to +40), with the top-average BLISS score and the highest-synergy 3×3 region (black outline) indicated. **D,** As in C, for 3D organoid models treated with RASi and AXLi (1500-6000 nM) for 5 days.

Building on these mechanistic insights, we next evaluated whether the efficacy of RAS inhibition could be enhanced by concurrent AXL inhibition. To address this possibility, we assessed cell proliferation following five days of treatment with the combination of AXLi and RASi or ERKi. Combined treatment with AXLi resulted in synergistic growth suppression (Bliss synergy score > 0) with RASi in nine of 10 cell lines and with ERKi in eight of 10 cell lines (Fig. 3C; Supplementary Fig. S3A and S3B). Similar results were also seen in 14-day clonogenic growth assays (Supplementary Fig. S3C and S3D) and in three-dimensional (3D) nonadherent spheroid growth assays (Supplementary Fig. S3E and S3F). Concurrent UNC2025 and daraxonrasib treatment also caused synergistic growth suppression (Supplementary Fig. S3G). These results provide further support for concurrent RASi and AXLi treatments.

To further validate our findings in preclinical models of PDAC that may better model patient response, we evaluated the combination treatment in patient-derived *KRAS*-mutant PDAC organoid 3D cell culture models that more accurately recapitulate tumor heterogeneity and three-dimensional proliferation within an extracellular matrix environment (47). Consistent with our short- and long-term treatment results in 2D monolayer and 3D cell line cultures, co-targeting AXL strongly and synergistically potentiated the growth-suppressive effects of RAS-ERK pathway inhibition in three organoid models with variable results in two others (Fig. 3D; Supplementary Fig. S3H and S3I).

To determine a mechanistic basis for the synergistic growth suppression caused by concurrent inhibition of AXL and RAS, we performed reverse phase protein array (RPPA) analysis to profile inhibitor-mediated changes in signaling activity. We treated a panel of four PDAC cell lines with RASi or AXLi alone or in combination at acute (4 hours), intermediate (24 hours), and prolonged (72 hours) time points (Supplementary Fig. S3J). While cell line variations were seen, some general trends were identified across three or four cell lines. At 4 hours, RASi treatment induced strong target inhibition, as indicated by downmodulation of pERK (T202/Y204) across all cell lines. In contrast, AXL inhibition caused little or no reduction in pERK. Further downstream, AXLi or RASi variably reduced phosphorylation of RPS6 at its ERK phosphorylation sites (S235/236) and its mTORC1-S6K phosphorylation sites (S240/244) that were further suppressed by combination treatment. Beginning at 24 hours and advancing further by 72 hours, rebound of pERK but not pRPS6 was observed in two of four cell lines.

KRAS promotes *MYC* gene transcription and MYC protein stability through phosphorylation by ERK and AKT (48). At 4 hours, RASi strongly reduced total MYC in three of four cell lines, but not in the RAS inhibitor−resistant KP-4 cells. In contrast, AXLi alone caused limited MYC reduction that was further enhanced by the combination. By 72 hours, partial reduction in MYC was maintained, with greater reduction upon combination treatment.

Beginning at 24 hours (Supplementary Fig. S3J), RASi, and to a lesser degree AXLi, reduced inhibitory phosphorylation of the RB tumor suppressor (S807/811), with stronger reduction seen with the combination. Similarly, each inhibitor alone caused limited reduction in expression of the anti-apoptotic protein Survivin (*BIRC5*), with the combination causing further reduction. Thus, KRAS and AXL inhibition cooperated to more effectively suppress cell cycle progression and cell survival.

AXLi treatment caused additional cellular responses more strongly than did RASi. At 24 hours, it robustly induced DNA damage, as indicated by increased phosphorylation of the double strand break marker γH2AX (S139). Additionally, AXLi strongly suppressed autophagy, as monitored by increased LC3B expression. Taken together, the enhanced growth suppression caused by the combination may be attributed to induction of both distinct and convergent cellular alterations (Supplementary Fig. S3K).

### AXL Drives RAS Inhibitor Resistance in *KRAS*-Mutant Lung Adenocarcinoma

We next determined if AXL could drive RAS inhibitor resistance in another cancer type with a high frequency of *KRAS* mutations. For these analyses we focused on the most prevalent subtype of lung cancer, LUAD, 30% of which harbors a *KRAS* mutation, predominantly the smoking-associated G12C (49,50). We utilized the KRAS^G12C^ inhibitor−sensitive LUAD cell line NCI-H358 (51–53) to determine if AXL overexpression alone is sufficient to drive resistance to RAS inhibition.

We stably infected NCI-H358 cells with lentivirus expression vectors encoding control green fluorescent protein (designated H358-GFP) or FLAG-tagged AXL (H358-AXL). Western blot analyses verified high steady-state AXL expression in H358-AXL cells (Supplementary Fig. S4A). Compared with H358-GFP cells, H358-AXL cells exhibited resistance (164- and 847-fold higher GI_50_, respectively) to the approved G12C-selective OFF-state inhibitor adagrasib and the RAS(ON) G12C-selective tri-complex inhibitor RMC-4998 (53) (Fig. 4A; Supplementary Fig. S4B). Resistance (31-fold higher GI_50_) to RASi was also observed. Thus, AXL overexpression alone was sufficient to drive strong resistance to treatment with both mutant-selective and RAS(ON) multi-selective RAS inhibitors.

**Figure 4.**
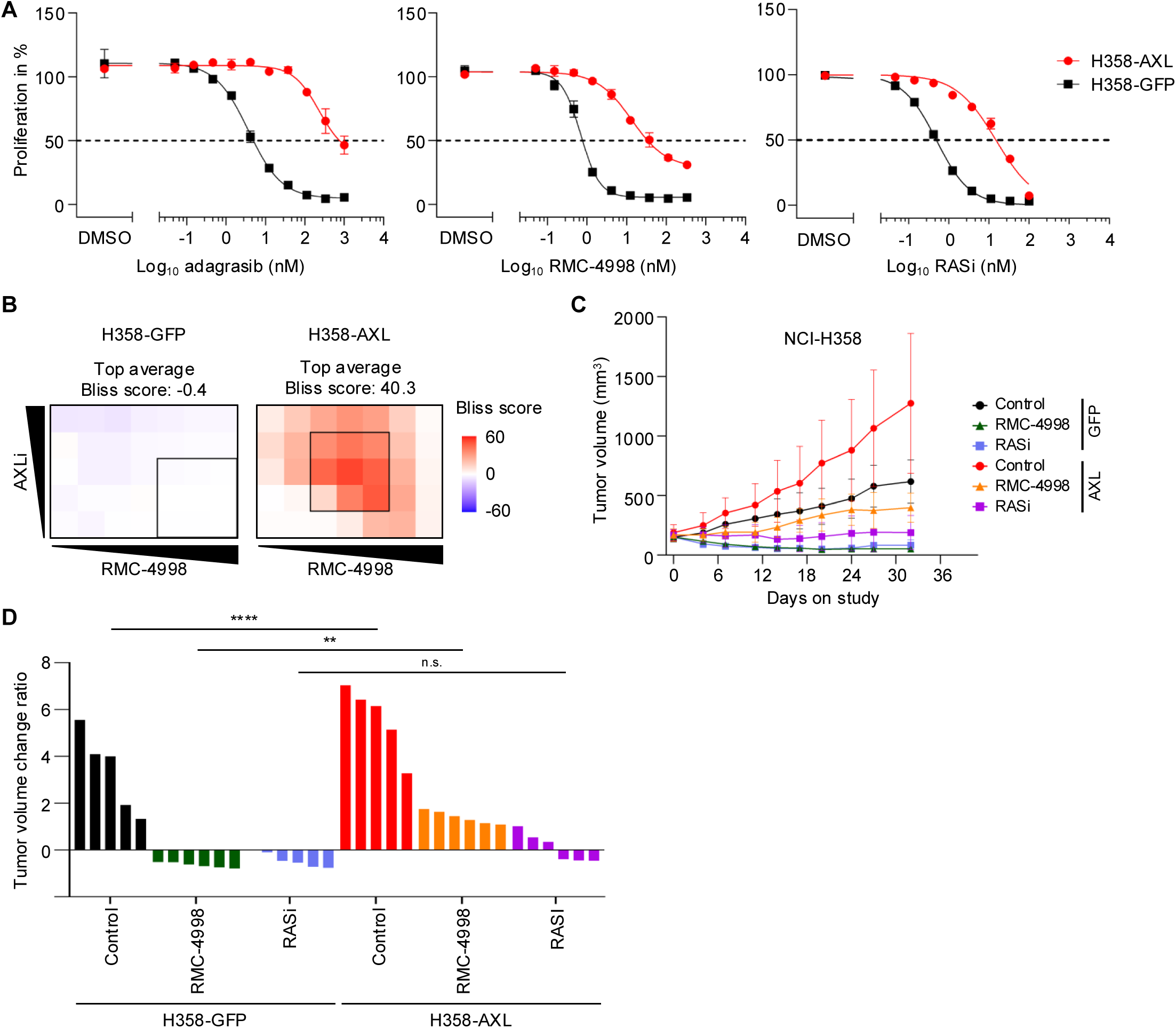
AXL overexpression confers resistance to RAS inhibition and creates a synergistic dependency on combined AXL inhibition. **A,** Proliferation of isogenic NCI-H358 cell lines ectopically expressing GFP (H358-GFP, black) or AXL (H358-AXL, red) treated with increasing concentrations of adagrasib, the RAS(ON) G12C-selective tricomplex inhibitor RMC-4998, or RASi for 5 days, shown as percent proliferation relative to DMSO; the dashed line indicates GI_50_. Points are the mean ± SEM (*n* = 3 biological replicates). **B,** BLISS synergy heatmaps for combined AXLi and RMC-4998 treatment in H358-GFP and H358-AXL cells grown in 3D spheroid culture. Color scale −60 to +60 (red, synergy; blue, antagonism). The highest-synergy region is outlined. **C,** Tumor growth of H358-GFP and H358-AXL xenografts treated with vehicle (control), G12Ci, or RASi; tumor volume (mm³) versus days on study (mean ± SD; *n* = 5-6 tumors per condition). **D,** Waterfall plot showing the relative change in individual tumor volume from day 0 to day 32 for H358-GFP and H358-AXL xenografts across the control, G12Ci, and RASi treatment groups. Values < 0 indicate tumor shrinkage. Statistical significance between treatment groups was determined by two-way ANOVA followed by Tukey’s multiple comparison test, (n.s. = not significant, \*\**P <*0.01, \*\*\*\**P <* 0.0001). All other comparisons are listed in Supplementary Fig. S4F.

We next determined whether concurrent AXL inhibition would enhance or restore sensitivity to KRAS^G12C^ inhibition. We found that, in 3D spheroid cultures, RMC-4998 inhibited the proliferation of H358-GFP cells, but this was not significantly enhanced by co-treatment with AXLi (Fig. 4B). In contrast, concurrent treatment with RMC-4998 and AXLi synergistically suppressed the proliferation of H358-AXL cells.

We then extended these analyses to *in vivo* models using cell line−derived xenograft (CDX) tumors established subcutaneously with H358-AXL or H358-GFP cells. Immunohistochemistry (IHC) staining confirmed AXL overexpression in H358-AXL tumors *in vivo* (Supplementary Fig. S4C). Tumor growth was monitored for 32 days under treatments with vehicle control, G12Ci (100 mg/kg orally once daily) or RASi (10 mg/kg orally once daily). Interestingly, although AXL overexpression did not significantly alter the adherent growth rate *in vitro* (Supplementary Fig. S4D), the tumorigenic growth of AXL-overexpressing CDX tumors in immune-compromised mice was greatly accelerated (Fig. 4C). All treatments were well tolerated as assessed by body weight throughout the study (Supplementary Fig. S4E). Whereas G12Ci induced tumor regression in H358-GFP tumors, only partial inhibition of tumor growth was observed in H358-AXL tumors (Fig. 4C and 4D; Supplementary Fig. S4F). Similarly, RASi led to tumor regression in H358-GFP tumors but only resulted in tumor stasis in H358-AXL tumors. Thus, AXL overexpression alone conferred partial resistance to RAS inhibitors *in vivo*. In summary, these findings demonstrated that AXL overexpression can drive resistance to RAS inhibitors in both PDAC and LUAD tumors, and that co-treatment with AXLi can restore RAS inhibitor sensitivity.

### Combined AXL and RAS Inhibition Exhibits Strong Synergistic Effects in RAS Inhibitor-Sensitive and -Resistant Cell Line-Derived Xenograft Tumors

To further assess the efficacy of combined AXL and RAS inhibition, we employed two human PDAC CDX tumor models in immune-compromised mouse models. Previously, we found that HPAF-II tumors exhibited robust regression when treated daily with 10 mg/kg RASi (14). To assess the synergistic potential of combination therapy, we utilized a suboptimal dose of RASi (3 mg/kg once daily) alone or in combination with AXLi (50 mg/kg twice daily) at a concentration shown previously to overcome PDAC tumor resistance to gemcitabine when used in combination (43). Under these experimental conditions, RASi but not AXLi alone caused a statistically significant but relatively limited impairment in tumor progression (Fig. 5A and 5B). Strikingly, combined AXLi and RASi treatment caused a significant synergistic effect. Furthermore, the tumor-inhibitory effect persisted for a further three days following treatment cessation (day 15) before the tumors resumed growth, resulting in an overall prolonged survival benefit for treated mice (Fig. 5C). In contrast to the single agents alone, the combination (comb.) increased median survival by nearly 3-fold, from 12.5 to 34.5 days (Supplementary Fig. S5A). The drug combination was well tolerated, with body weight remaining stable throughout treatment, and no evidence of liver toxicity was observed (Supplementary Fig. S5B and S5C).

**Figure 5.**
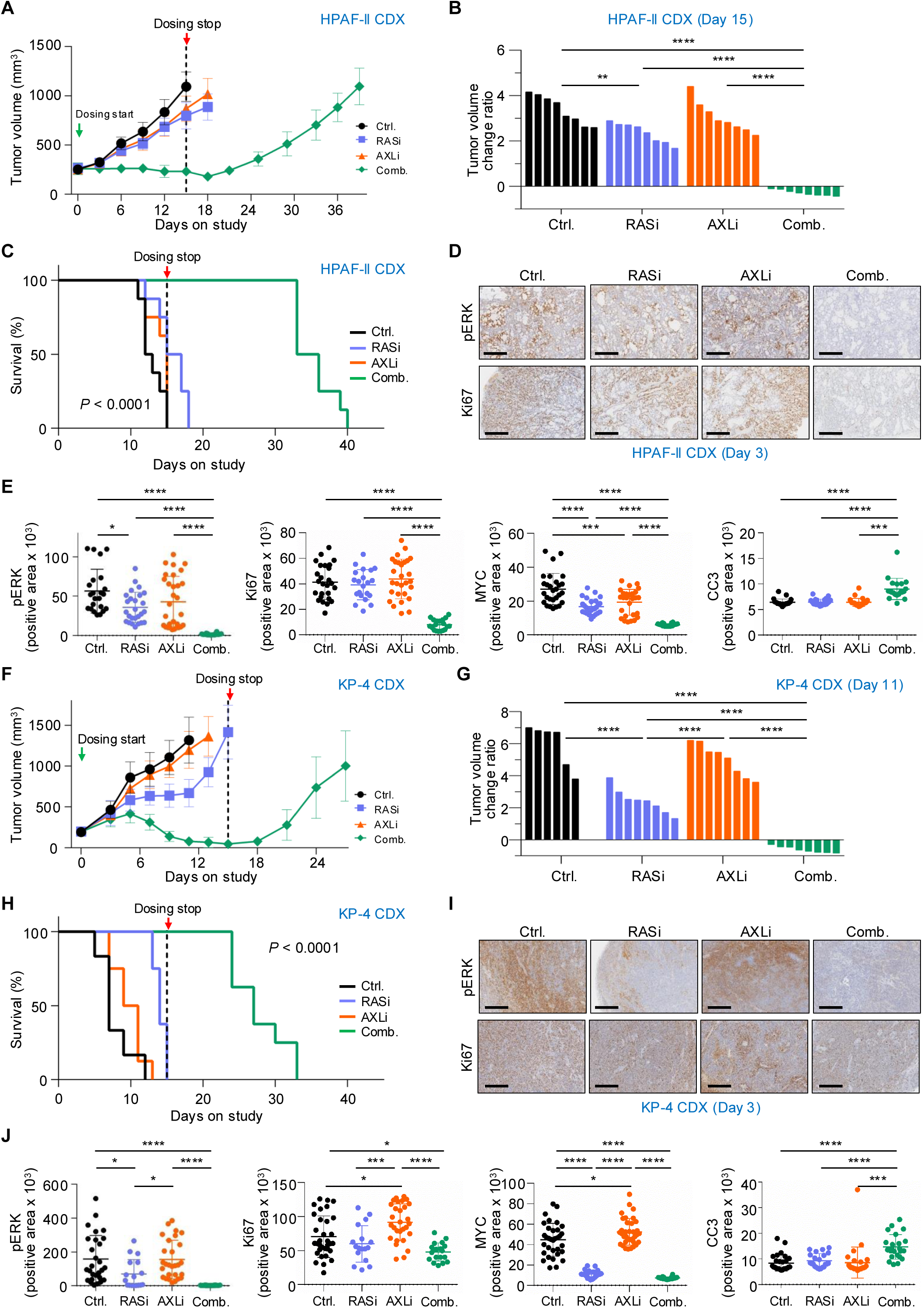
AXLi enhances RASi antitumor efficacy in RASi-sensitive and -resistant PDAC CDX models. **A,** Tumor volume plotted over time for RASi sensitive HPAF-II tumors treated for 15 days with vehicle (black), RASi (blue), AXLi (orange), or the combination of RASi and AXLi (green). The dashed vertical line indicates end of treatment at day 15. Data shown as mean ± SD, *n* = 8 tumors per condition. **B,** Waterfall plot of normalized individual tumor volume changes day 0 versus day 15 (end of treatment) of HPAF-II tumors across the control, RASi, AXLi and combination treatments. Values < 0 indicate tumor shrinkage. Statistical significance was determined by one-way ANOVA followed by Tukey’s multiple comparison test (\*\**P* <0.01, \*\*\*\**P <* 0.0001). Only statistically significant results are shown. **C,** Kaplan-Meier survival curve of HPAF-II tumor-bearing mice treated as indicated. Differences among treatment groups were assessed using a global log-rank (Mantel–Cox) test; \*\*\*\**P <* 0.0001. **D,** Representative immunohistochemical staining for pERK and Ki67 in HPAF-II tumor sections. Scale bars indicate 600 µm. **E,** Quantitative analysis of staining for pERK, Ki67, MYC, and cleaved caspase-3 (CC3). Each dot represents an analyzed area from an individual image under the indicated conditions. Data shown as mean ± SD. Statistical significance was determined using one-way ANOVA followed by Tukey’s multiple comparison test; \**P <* 0.05, \*\*\**P <* 0.001, \*\*\*\**P <* 0.0001. **F,** Tumor volume plotted over time for RASi-resistant KP-4 tumors treated for 15 days with vehicle (black), RASi (blue), AXLi (orange), or the combination of RASi and AXLi (green). (*n* = 6 - 8 tumors per condition). **G,** Waterfall plot showing the relative change in individual tumor volume from day 0 to day 11 (when the control group was sacrificed) of KP-4 xenografts across the control, RASi, AXLi and the combination treatment groups. Values < 0 indicate tumor shrinkage. Statistical significance was determined by one-way ANOVA followed by Tukey’s multiple comparison test (\*\*\*\**P <* 0.0001). Only statistically significant results are shown. **H,** Kaplan-Meier survival curve of KP-4 tumor-bearing mice treated as indicated. Differences among treatment groups were assessed using a global log-rank (Mantel-Cox) test; \*\*\*\**P <* 0.0001. **I,** Representative immunohistochemical staining for pERK and Ki67 in KP-4 tumor sections. Scale bars indicate 600 µm. **J,** Quantitative analysis of staining for pERK, Ki67, MYC, and CC3. Each dot represents an analyzed area from an individual image under the indicated conditions. Data shown as mean ± SD. Statistical significance was determined using one-way ANOVA followed by Tukey’s multiple comparison test; \**P <* 0.05, \*\*\*\**P <* 0.0001.

Consistent with the lack of anti-tumor activity, IHC analysis of tumors collected after three days of single agent treatment revealed minimal reductions in pERK and total MYC (Fig. 5D and 5E). In contrast, the combination treatment resulted in pronounced suppression of these markers of target inhibition, as well the proliferation marker Ki67 and the mitosis marker phosphorylated histone H3 (pHH3) (Fig. 5D and 5E; Supplementary Fig. S5D and S5E). Additionally, staining for the apoptotic marker cleaved caspase-3 (CC3) indicated enhanced apoptotic cell death in tumors receiving combination but not monotherapy treatment.

We next evaluated the combination therapy in the RAS inhibitor-resistant PDAC cell line KP-4 (36,54). KP-4 CDX tumors reached maximum volume by day 11, underscoring their aggressive growth phenotype (Fig. 5F and 5G) and requiring termination at that time. Like our observations in the RASi-sensitive HPAF-II CDX model, AXLi alone had no effect on KP-4 tumor growth. However, in the RASi-resistant KP-4 CDX model, whereas an optimal concentration of RASi alone (8) (10 mg/kg once daily) caused a limited tumor growth delay, the combination caused a striking degree of tumor shrinkage and furthermore delayed tumor regrowth by approximately seven days following the cessation of therapy on day 15 (Fig. 5F). Moreover, adding AXLi to RASi produced a near doubling of median overall survival compared to RASi alone (27 vs 14 days) (Fig. 5H; Supplementary Fig. S5F). As in the HPAF-II CDX model, no substantial changes in body weight or signs of liver toxicity were observed with any treatment of KP-4 tumor-bearing mice (Supplementary Fig. S5G and S5H).

IHC analysis showed that, while AXLi treatment alone had little to no impact on pERK or growth (Ki67 and pHH3), these were strongly reduced by the combination of AXLi and RASi (Fig. 5I and 5J; Supplementary Fig. S5I and S5J). The combination also strongly suppressed MYC expression and enhanced apoptosis (CC3-positive cells). Thus, in both CDX models, the combination caused significant suppression of key KRAS effector signaling activity.

### Combined RAS and AXL Inhibition Induces Tumor Regression and Prolongs Survival in Immunocompetent PDAC Models

To determine the impact of the combined KRAS and AXL inhibition in mouse PDAC models with a fully competent immune system, subcutaneous or orthotopic allograft tumors were established using the mouse PDAC cell line FC1242 derived from KPC (*Kras*^G12D^; *Trp53*^R172H^; *Pdx-Cre*) mouse tumors (55) backcrossed to the C57BL/6 genetic background (56). Subcutaneous versus orthotopic PDAC tumors are distinct in their tumor microenvironment regarding stromal desmoplasia content and vascularity, with the latter providing a more clinically relevant setting (57). However, subcutaneous models have the experimental advantages of improved drug delivery and ease of monitoring tumor size. Tumor-bearing mice were treated for 15 days with vehicle control, low dose RASi (3 mg/kg once daily), AXLi (50 mg/kg twice daily), or the combination.

In the subcutaneous allografts, whereas AXLi alone did not significantly impair tumor growth and RASi alone produced a 50% decrease in tumor growth relative to vehicle-treated controls, the combined inhibition of RAS and AXL induced objective tumor regressions in all treated animals (Fig. 6A and 6B). The combination also conferred a marked survival advantage, with a median survival of 42 days despite cessation of dosing after day 15, and doubling the median survival compared to RASi alone (Fig. 6C and 6D). Little change in weight was observed in any of the tumor-bearing animals (Supplementary Fig. S6A), indicating that the treatments were well tolerated.

**Figure 6.**
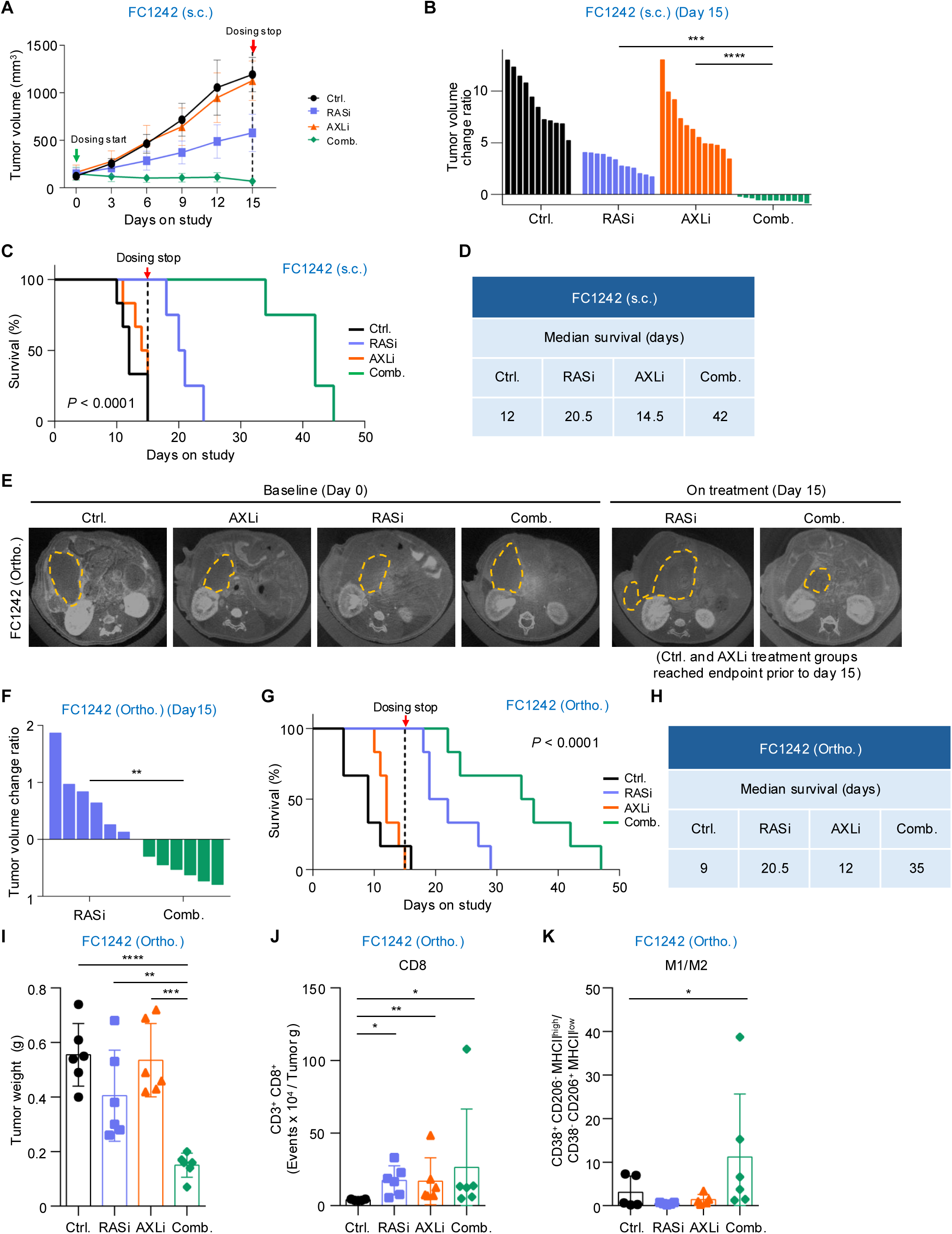
Combined AXLi and RASi treatment extends survival and remodels the immune microenvironment in syngeneic PDAC models. **A-D**, subcutaneous (s.c.) implantation of FC1242 PDAC cells; **E-K**, orthotopic (ortho.) implantation of FC1242 PDAC cells. **A,** Tumor growth of subcutaneous (s.c.) FC1242 cell line−derived allografts (KPC-derived *Kras*^G12D^; *Trp53*^R172H^ murine PDAC, implanted in immunocompetent F1 C57Bl/6 J x 129S2/Sv mice treated with vehicle (Ctrl.), RASi, AXLi or the combination. Tumor volume (mm³) with dosing start and stop indicated (mean ± SD; *n* = 11-12 tumors per group). **B,** Waterfall plot showing the relative change in individual tumor volume from day 0 to day 15 (*n* = 6 tumors per group). Statistical significance determined using one-way ANOVA followed by Sidak’s multiple comparison test (\*\*\**P <* 0.001, \*\*\*\**P <* 0.0001). All other comparisons are listed in Supplementary Fig. S6J. **C,** Kaplan-Meier survival curve of subcutaneous FC1242-bearing mice treated as indicated; the dashed line indicates dosing stop. Differences among treatment groups were assessed using a global log-rank (Mantel–Cox) test; \*\*\*\**P <* 0.0001. **D,** Summary of the median survival (days) for the indicated treatment groups in FC1242 subcutaneous model shown in C. **E,** Representative microCT images of orthotopic (ortho.) FC1242 tumors (outlined by dashed lines) at baseline (day 0) and on treatment (day 15). Control and AXLi groups reached endpoint before day 15. **F,** Waterfall plot showing the relative change in individual tumor volume from day 0 to day 15 (end of treatment) for the FC1242 orthotopic model (*n* = 6 tumors per group). Statistical significance was determined using unpaired t-test with Welch’s correction (\**\*P <* 0.01). **G,** Kaplan-Meier survival curve of orthotopic FC1242-bearing mice treated as indicated (*n* = 6 per group). Differences among treatment groups were assessed using a global log-rank (Mantel-Cox) test; \*\*\*\**P <*0.0001. **H,** Summary of the median survival (days) for the FC1242 orthotopic model shown in **G. I,** Tumor weights in g of orthotopic FC1242 PDAC tumors from the immunophenotyping cohort collected at day 10 of indicated treatment groups (n = 6 tumors per group). Statistical significance was determined using one-way ANOVA followed by Tukey’s multiple comparison test (\*\**P* < 0.01, \*\*\**P* < 0.001, \*\*\*\**P* < 0.0001). **J**, Intratumoral CD8⁺ T-cell density (CD3⁺CD8⁺ events × 10⁴ per gram of tumor). **K,** Tumor macrophage polarization, expressed as the M1/M2 ratio (CD38⁺CD206⁻MHCII^high / CD38⁻CD206⁺MHCII^low). **J-K,** Data shown as mean ± SD. Comparisons were made by Kruskal–Wallis test with FDR correction using the two-stage step-up method of Benjamini, Krieger and Yekutieli (*q < 0.05, **q < 0.01). Only significant comparisons are shown.

In the orthotopic setting, whereas tumors in the RASi and combination groups remained evaluable by day 15, all animals in the vehicle and AXLi monotherapy groups had perished due to tumor burden. MicroCT imaging at day 15 demonstrated reduced tumor burden upon RASi treatment, with tumor regression observed in the combination group (Fig. 6E). As we observed in the subcutaneous setting, the combination caused significantly greater tumor reduction compared with RASi alone (Fig. 6F). Furthermore, while RASi alone substantially increased median survival (20.5 days) compared to vehicle (9 days) or AXLi 12 days), the combination conferred a robust additional extension (to 35 days; Fig. 6G and 6H). Little change in weight was observed in any of the tumor-bearing animals (Supplementary Fig. S6A), indicating that the treatments were well tolerated. Together, these data demonstrated that combined targeting of RAS and AXL in immunocompetent PDAC models produces deeper and more durable anti-tumor responses than either inhibitor alone, closely mirroring our observations with the two human CDX models.

To assess whether the combination treatment elicited changes in the tumor microenvironment associated with an enhanced antitumor response, we performed multiparameter flow cytometry on orthotopic tumors following 10 days of treatment. Because animals in our survival studies reached endpoint rapidly once tumors were large, immunophenotyping was initiated at a baseline tumor volume of 100-120 mm³ (i.e., day 10). At day 10, only the combination caused significant tumor reduction (Supplementary Fig. S6B) This timing ensured sufficient treatment exposure and permitted tissue collection before premature mortality. To account for differences in tumor size at the time of collection (Fig. 6I), immune populations were normalized to tumor mass.

Suboptimal RAS inhibition alone significantly increased intratumoral CD8+ T-cell abundance (Fig. 6J), consistent with prior reports describing immune alterations following Kras inhibition with optimal doses of RASi (58,59). AXLi alone also significantly increased CD8+ T cell frequency (Fig. 6J). RASi alone caused limited enhancement in the frequency of NK and NK-T cells while AXLi enhanced only NK-T cells (Supplementary Fig. S6D and S6E). Yet the combination further enhanced NK but not NK-T cells.

However, the combination did not further increase lymphoid populations beyond the effect of single inhibition of RAS or AXL (Fig. 6J; Supplementary Fig. S6F-S6H). Analysis of the myeloid compartment revealed a higher frequency of the CD11b^+^ Ly6G^−^ resident monocyte and the F4/80^+^ macrophage populations in both the AXLi and combination arms (Supplementary Fig. S6I and S6J), suggesting specific AXLi-driven changes. However, there were no differences in these between AXLi alone and combination treatments. Instead, concurrent inhibition of RAS and AXL produced a distinct shift in macrophage polarization, resulting in an increased M1/M2 ratio relative to both vehicle and single-agent groups (Fig. 6K). This shift toward an M1-like phenotype is consistent with enhanced pro-inflammatory macrophage activity. No consistent combination-specific changes were observed across other myeloid subsets examined (Supplementary Fig. S6K-S6M).

Together, these findings show that co-targeting RAS and AXL caused comparable tumor regression and durable survival benefit in both immunocompetent and immunodeficient mouse models of PDAC. While RASi and AXLi induced shared and distinct consequences on the immune environment, the superior efficacy of the combination is primarily associated with enhanced tumor-intrinsic suppression of RAS signaling.

### AXLi Potentiates RASi-driven Transcriptional Remodeling in CDX Tumors

To characterize the transcriptional consequences of AXLi, RASi, and their combination in tumors, we performed RNA-sequencing on HPAF-II and KP-4 CDX tumors following three days of treatment to capture the immediate consequences of target inhibition. Consistent with its limited impact on tumor growth, AXLi monotherapy produced only negligible transcriptomic changes in HPAF-II and KP-4 tumors, with fewer than 10 significantly dysregulated genes detected in either model (Fig. 7A and 7B). By contrast, RASi elicited a substantially broader transcriptional response, including both induced (upregulated; UP) and suppressed (downregulated; DN) transcripts. A suboptimal dose of RASi caused relatively few changes in gene transcription in HPAF-II tumors (262 UP/DN genes) but had a far more pronounced effect in KP-4 tumors (3,310 UP/DN genes). Combined treatment with RASi and AXLi further expanded the set of differentially expressed genes, yielding an approximately 1.5-fold increase in KP-4 tumors (4,816 UP/DN genes) and a nearly 20-fold increase in HPAF-II tumors (5,133 UP/DN genes) relative to RASi monotherapy.

**Figure 7.**
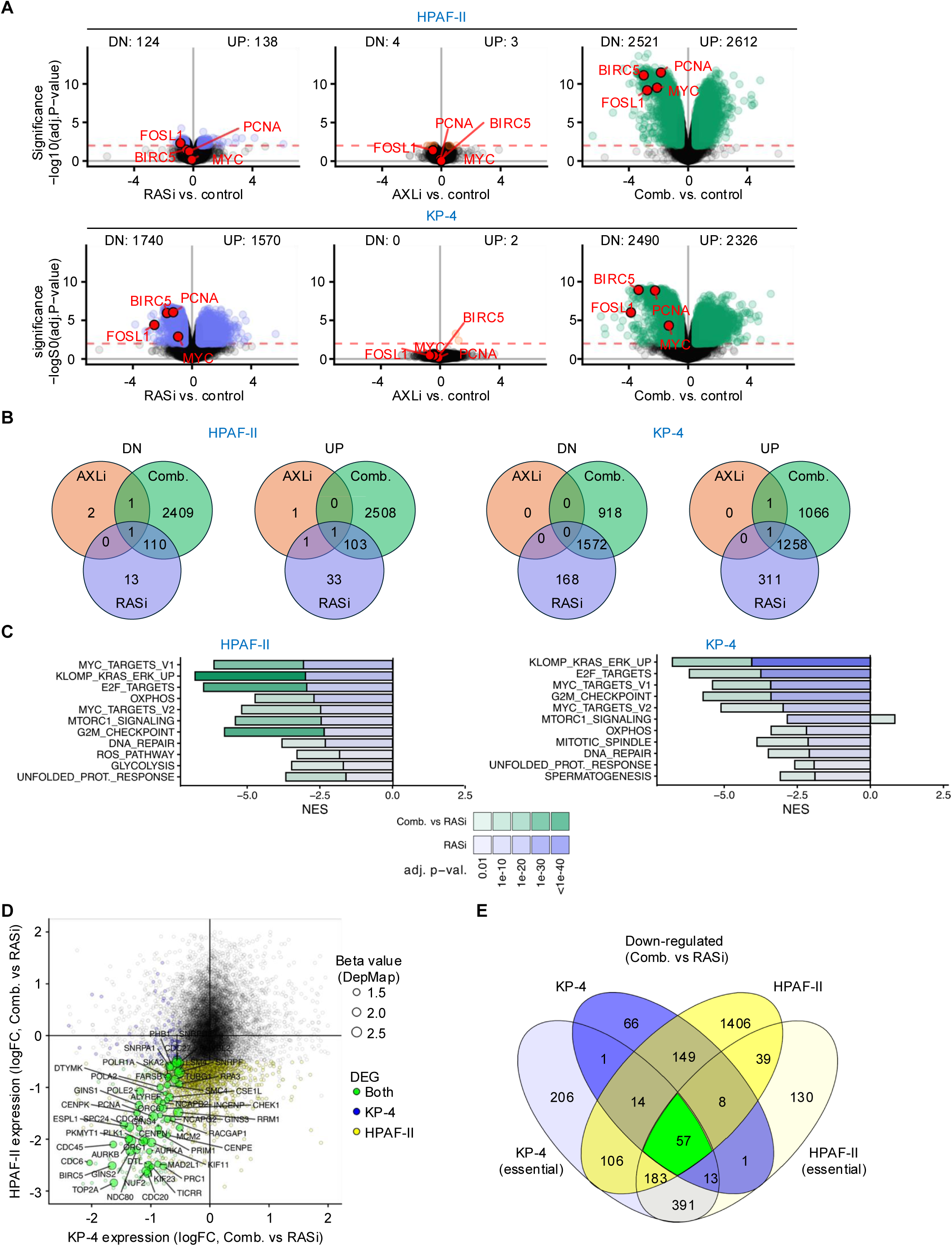
Combination therapy with RASi and AXLi enhances RASi-dependent transcriptional changes in both RASi-sensitive and RASi-resistant CDX models. **A,** Volcano plots of differentially expressed genes (shaded circles) in RASi-sensitive HPAF-II (top) and RASi-resistant KP-4 (bottom) tumors harvested on day 3 of treatment, comparing RASi versus control, AXLi versus control (Ctrl.), and combination (Comb.) versus control. Numbers of significantly up- and downregulated genes are indicated above each plot with a significance threshold of FDR < 0.01 (dashed red line) and logFC > 0.5. Representative KRAS-dependent genes indicated in red. *n* ≥ 4 for each group. **B,** Venn diagrams showing the overlap of differentially expressed genes among RASi-, AXLi-, and combination-treated tumors in A, shown separately for downregulated and upregulated genes in HPAF-II (left panels) and KP-4 models (right panels) (*n* ≥ 4 tumors per condition). **C,** Gene set enrichment analysis (GSEA) in HPAF-II (left) and KP-4 (right) xenografts harvested on day 3 of treatment. For each gene set, stacked bars show the normalized enrichment score (NES) for RASi versus control (blue) and the additional contribution of AXLi (Comb. versus RASi, green). The gene sets comprise Hallmark programs together with a curated KRAS-ERK signature (KLOMP_KRAS_ERK_UP) (9). All depicted gene sets are negatively enriched in RASi treated tumors and further negatively enriched in Comb. treated tumors (except KP-4 MTORC1_SIGNALING). Bar color intensity reflects the adjusted P value (darker, more significant). *n* ≥ 4 tumors per condition. **D,** Scatterplot comparing the transcriptional effects of the combination over RASi (logFC, Comb. vs RASi) in KP-4 (x-axis) versus HPAF-II (y-axis). Each point is a gene. Differentially expressed genes (DEGs) are colored by model (KP-4 only, blue; HPAF-II only, yellow; both models, green). Point size reflects the gene-dependency beta value from DepMap (larger = stronger dependency). **E,** Venn diagram of genes downregulated by the combination relative to RASi (Comb. vs RASi) across KP-4 and HPAF-II models and their intersection with essential genes (KP-4 essential, HPAF-II essential). Numbers indicate genes in each overlap. The central intersection (green, *n* = 57) comprises genes downregulated only by the combination vs RASi alone in both models that are also essential in both, pointing at a core set of genetic dependencies that are suppressed by the combination but not by RASi alone.

That concurrent AXLi enhanced the transcriptional changes seen with RASi alone was recapitulated in selected markers representing major ERK-driven transcriptional and phenotypic outputs, including *MYC* and *FOSL1* (FRA1, a component of the ERK-responsive transcription factor AP-1), *PCNA*, and, consistent with our RPPA findings, *BIRC5* (Survivin) (Fig. 7A; Supplementary Fig. S7A). Summarization of these differentially expressed genes reinforced that AXLi monotherapy contributed very few uniquely regulated transcripts in either model, whereas combined AXLi and RASi treatment markedly expanded both the RASi UP and DN gene sets (Fig. 7A and 7B). In both KP-4 and HPAF-II tumors, combination inhibitor treatment resulted in changes in expression of genes that are predominantly KRAS-dependent in PDAC (9). In KP-4 tumors, the combination response partially built on the transcriptional program elicited by RASi alone, with substantial overlap between the two treatment arms. In contrast, HPAF-II tumors displayed a predominance of combination-specific transcripts, i.e., the 2521 DN and 2612 UP genes indicate that AXLi substantially enhances the transcriptional response to suboptimal RASi in this model.

Given the minimal transcriptional effects of AXLi monotherapy, we next sought to define its specific contribution in the combination setting. There, AXLi increased both the number of RASi-responsive genes and the magnitude of their induction or repression, indicating that its primary effect was not to initiate an independent transcriptional program but rather to deepen the response elicited by RASi. Consistent with this, gene set enrichment analysis (GSEA) showed stronger suppression of the KRAS_ERK_UP (9), MYC_UP (74), and Hallmark_E2F Targets (60) programs upon combined AXLi and RASi treatment than upon RASi treatment alone (Fig. 7C). More broadly, pathway-level analysis with Hallmark gene sets confirmed that KRAS-/ERK-, MYC-, and E2F-associated gene sets were among the most strongly suppressed pathways in both tumor models, and this effect was still evident in HPAF-II tumors despite incomplete pathway suppression by RASi monotherapy (Fig. 7C; Supplementary Fig. S7B). Moreover, that all depicted gene sets were negatively enriched indicates that RASi treatment resulted in coordinated downregulation across well-established RAS-dependent proliferative (e.g., MYC) and metabolic (e.g., OXPHOS) transcriptional programs (9), with the combination deepening this suppression.

The key effect of AXLi in the presence of RASi treatment was to substantially further reduce expression of the oncogenic Hallmark programs E2F_TARGETS, MYC_TARGETS_V1, and G2M_CHECKPOINT in both tumor models, and this was most evident in the HPAF-II CDX model (Fig. 7C). In both cases, the most robust set of gene expression changes upon single agent RASi or combination treatment was our previously identified RAS-ERK dependent signature, KLOMP_KRAS_ERK_UP (9).

To determine whether adding AXLi to RASi results in the regulation of further genes that are essential for cellular proliferation and tumor growth, we compared the differentially expressed transcripts resulting from combination treatment versus RASi monotherapy (Fig. 7D). Upon assigning the AXL-dependent genes shared between the two cell lines to their Cancer Dependency Map (DepMap) CRISPR beta values, we noticed that a large set of essential genes required for cell cycle progression was downregulated in both models (Supplementary Fig. S7C). A comparison of these sets identified 57 genes that were both essential and downregulated by the addition of AXLi to RASi treatment in both HPAF-II and KP-4 (Fig. 7E, green), and another 183 that were essential in both tumor models but downregulated upon addition of AXLi only in the HPAF-II tumor model (Fig. 7E), likely due to the stronger RASi monotherapy effect seen in the KP-4 model (Fig. 5A and 5F). Together, these data support an unanticipated mechanism wherein RAS inhibition induces GAS6 expression and subsequent activation of AXL-mediated antagonism of the efficacy of RAS inhibitors to block KRAS function (Supplementary Fig. S7D). Therefore, concurrent treatment with an inhibitor of AXL kinase function enhances the ability of RAS inhibitors to suppress KRAS effector signaling, and this explains how the combination promotes synergistic anti-tumor activity.

## DISCUSSION

The recent observations from phase 3 clinical evaluation of the RASi daraxonrasib support the therapeutic value of anti-RAS therapies for PDAC (16). The OS of patients who received daraxonrasib was a striking 13.2 months versus 6.7 months for patients who received chemotherapy. However, with an objective response rate of 32%, primary as well as acquired resistance (61,62) limit the depth and durability of daraxonrasib clinical efficacy. To overcome these limitations, combination therapy emerges as a promising strategy to enhance the clinical efficacy of RAS inhibitors in general. While the appearance of DNA alterations in initially responsive patients who relapse on daraxonrasib monotherapy provides one guide for the development of useful combinations, in ∼50% of such patients, no DNA alterations were identified (61,62). To identify nongenetic mechanisms of resistance to RAS-ERK inhibitors, we applied an oncology drug chemical biology screen and a druggable genome CRISPR genetic screen and identified the AXL receptor tyrosine kinase as a putative resistance driver. Unlike other oncogenic RTKs, AXL activation in cancer involves nongenetic mechanisms (41,63). Here we determined that AXL overexpression is sufficient to drive RAS-ERK inhibitor resistance and that AXL inhibition synergistically enhanced RAS inhibitor antitumor activity to cause regressions in preclinical PDAC and LUAD tumor models. Independently, another study applied a gain-of-function screen and identified AXL as a driver of resistance to the KRAS^G12D^-selective inhibitor MRTX1133 (64). That study also showed that concurrent treatment with the AXLi bemcentinib and MRTX1133 caused synergistic tumorigenic growth suppression in mouse models of lung cancer. Thus, it is likely that this class of combination therapy may be applicable broadly to other RAS inhibitors in other RAS-mutant cancer types.

Addressing a molecular basis for the synergistic activity of this combination, we identified an unexpected mechanism wherein AXL attenuated the ability of a RAS inhibitor to suppress effector signaling through ERK and MYC. Consistent with this finding, but inconsistent with currently known mechanisms of AXL signaling (41,63), we observed that concurrent pharmacological inhibition of AXL and RAS synergistically enhanced suppression of ERK and MYC. These findings provide a molecular explanation for the ability of the combination to drive persistent tumor regression and an enhanced survival advantage preclinically. Together, they support the clinical evaluation of combinations with AXL inhibition as a therapeutic strategy to enhance anti-RAS therapies.

Our identification of AXL as a driver of resistance to RAS-ERK inhibitors extends previous observations implicating AXL in driving resistance to chemotherapy and other targeted therapies (25,63,65,66). For example, Brekken and colleagues showed that AXLi enhanced PDAC tumor response to gemcitabine (43). We previously applied a CRISPR screen and identified AXL as a driver of resistance to MEK and ERK inhibitors in PDAC cell lines (40), and we showed that AXLi potentiated the effect of MEKi in PDAC cell lines and patient-derived organoids (67). Similarly, AXL was identified as a RASi resistance mechanism in a CRISPR screen using the KRAS^G12C^-selective inhibitor ARS-1620 in the PDAC cell line MIA PaCa-2 (68) and increased AXL activation was associated with PDAC cell line resistance to MRTX1133 (36). We also found that elevated AXL expression was associated with YAP-TEAD-driven resistance to the FDA-approved KRAS^G12C^-selective inhibitor adagrasib (69).

Here, we validated the effectiveness of concurrent AXL and RAS inhibition as a therapeutic approach in cell line, organoid, and human and mouse tumor models of PDAC. Our *in vivo* analyses included two immunodeficient human PDAC CDXs and immunocompetent subcutaneous and orthotopic syngeneic mouse models. A consistent pattern was observed in all four models, where AXLi alone showed very limited or no antitumor activity, RASi alone reduced the rate of tumorigenic growth, and the combination of RASi and AXLi caused tumor regression.

Although we found that AXLi inhibited PDAC growth in cell line and organoid models, unexpectedly, AXLi monotherapy showed limited/no anti-tumor activity, suggesting that the tumor microenvironment may impact AXLi responsiveness. A similar lack of AXLi activity was also described in PDAC patient-derived xenograft subcutaneous tumor models (43). Despite this lack of monotherapy efficacy, when combined with RAS inhibition, we observed synergistic tumor regression in both immune-deficient and -competent mouse models. This finding is at odds with the prevailing mindset that effective combinations must be comprised of drugs that each alone exhibits anti-tumor efficacy. Thus, when transitioning our findings to clinical evaluation, the requirement to first establish AXLi monotherapy may not be a valid go/no-go requirement. Embracing this different mindset, we have initiated clinical evaluation of the combination of AXL and KRAS inhibition in PDAC.

Evaluation of the consequences of genetic or pharmacological suppression of AXL function has established a critical role for AXL in promoting an immune suppressive environment (23–25). However, our evaluation of the impact of drug treatment on immune cells in the orthotopic syngeneic mouse model found an unexpectedly limited role for an immune response in contributing to the synergistic activity seen with the combination of AXLi and RASi. Several immune populations were increased in the presence of single RASi (CD8+ T and NK cells) and AXLi (CD8+ T and NK-T cells, resident monocytes, macrophages and granulocytic MDSCs). Some of these results are consistent with previous studies reporting enhanced frequency of CD8+ T cells in response to single inhibition of either KRAS (58,59) or AXL (70–73). Nonetheless, only a modest shift to the M1 inflammatory state was observed exclusively under the combination treatment. This minimal remodeling of the tumor immune microenvironment by the combination aligns with the nearly identical anti-tumor activities seen in the immunodeficient and immunocompetent mouse models. We conclude that the synergistic antitumor activity of concurrent RAS and AXL inhibition is mediated predominantly through a tumor cell−autonomous mechanism through enhanced suppression of KRAS signaling.

Our IHC analyses in the CDX models showed that the degree of ERK and MYC suppression correlated with the degree of tumor suppression. Although AXL can activate RAS and ERK signaling, surprisingly, inhibition of AXL alone was not sufficient to significantly reduce either ERK activity or MYC expression. Instead, our observations are consistent with previous studies reporting that pharmacological combinations based on KRAS pathway inhibitors (i.e., of KRAS, MEK and ERK) required a profound degree of MYC suppression to obtain deeper, more durable anticancer responses in PDAC and LUAD (14,28,74–76).

Finally, we performed gene transcription analyses and identified an unexpected basis for the synergistic activity of the combination. First, although AXL is known to stimulate signaling networks (e.g., RAS-ERK) that should significantly impact gene transcription, surprisingly, inhibition of AXL alone caused essentially no significant transcriptional alterations. In contrast, and as expected, RASi alone caused significant transcriptional changes that were consistent with KRAS inhibition. More surprisingly, when RASi was combined with AXLi, the overall changes in gene transcription largely reflected a stronger suppression of KRAS-regulated gene transcription. Maitra and colleagues reached a similar conclusion in their studies in lung cancer (64). Together, these findings support a model in which, basally, there is no AXL activity in PDAC. The loss of ERK-MYC signaling upon RAS inhibition causes a compensatory feedback response that increases expression of the AXL ligand GAS6 and activation of AXL. This leads to AXL impairment of the efficacy of RASi, so when AXLi is combined with RASi, it restores or enhances the ability of RASi to suppress KRAS signaling and results in greater efficacy than either AXLi or RASi alone. This model proposes that AXL-mediated resistance to RASi is driven through a mechanism distinct from AXL downstream signaling. In summary, our study established a role for AXL in driving resistance to RAS inhibitors and supports the potential therapeutic value of concurrent AXL inhibition to enhance the efficacy of RAS inhibitors.

## METHODS

### Cell Lines

The human PDAC cell lines BxPC-3, CFPAC, HPAC, HPAF-II, MIA PaCa-2, KP-4, PANC-1 and SW1990 were obtained from American Type Culture Collection (ATCC). The patient-derived xenograft (PDX) human PDAC cell lines Pa01C, Pa14C, Pa16C, MDA-PATC50, MDA-PATC53, and MDA-PATC43 were provided by Dr. Anirban Maitra or Jason Fleming (MD Anderson Cancer Center), and UM53 was provided by Dr. Diane Simeone (University of Michigan). The mouse PDAC cell lines p53 2.1.1^syn-Luc^ and INK4.1^syn-Luc^ or FC1242 were provided by Dr. Eric Collisson (University of California, San Francisco) or Dr. David Tuveson (Cold Spring Harbor Laboratory), respectively. All cell lines were grown in a humidified chamber (37°C and 5% CO_2_) and maintained in either Dulbecco’s Modified Eagle Medium (DMEM) or Roswell Park Memorial Institute Medium (RPMI 1640), supplemented with 10% fetal bovine serum (FBS) and 1% penicillin/streptomycin, for no more than 15 passages. Cells were tested negative for Mycoplasma using MycoAlert Mycoplasma Detection Kit (Lonza) and authenticated by short tandem repeat (STR) analysis.

### Establishment of AXL Overexpressing Cells

NCI-H358 cells stably overexpressing control GFP or AXL were generated by infection with lentiviral expression vectors encoding MYC-DDK tagged human AXL (Origene Catalog #RC206431L3V) or mGFP (#PS100093V). Cells were selected and maintained in tissue culture under selective pressure with 2 µg/mL puromycin.

### Patient-Derived PDAC Organoids

The hM1A and hT105 patient-derived organoids were provided by Dr. David Tuveson (Cold Spring Harbor Laboratory) and have been described previously (77). The PT3, PT6 and PT8 KRAS^G12V^-mutant PDAC patient-derived organoids were provided by Dr. Calvin Kuo (Stanford) and have been described previously (78). Organoids were grown in growth factor reduced Matrigel (Corning) domes in complete human feeding medium in a humidified chamber (37°C and 5% CO_2_): The complete human medium was comprised of Advanced DMEM/F12 (Thermo Fisher Scientific) with WRN (L-WRN; ATCC CRL-3276) and supplemented with 500 nM A83-01 (TOCRIS), 1x B27 supplement (Thermo Fisher Scientific), 0.01 µM GlutaMAX (Thermo Fisher Scientific), 10 mM HEPES (Thermo Fisher Scientific), 50 ng/mL hEGF (Peprotech), 100 ng/mL hFGF10 (Peprotech), 0.01 µM hGastrin I (TOCRIS), 1.25 mM, 1 mM N-acetylcysteine (Sigma-Aldrich) and 10 mM nicotinamide (Sigma-Aldrich). For the first 2 days after reseeding as single cells, the growth medium was supplemented with 10.5 µM Y-27632 (Selleckchem) for all organoid models, and additionally with 10 µM SB202190 (Sigma-Aldrich). Organoids were tested routinely for mycoplasma using MycoAlert Mycoplasma Detection Kit (Lonza) and determined to be negative.

### Chemical Compounds

The RAS(ON) multi-selective tri-complex inhibitor RMC-7977 was provided by Revolution Medicines (8). The AXL selective inhibitor bemcentinib (BGB324) was obtained from Selleckchem. The MERTK inhibitor UNC2025 was provided by Dr. Xiaodong Wang (University of North Carolina at Chapel Hill). RMC-4998 and adagrasib were synthesized and characterized as previously described (51,53).

### Drug Sensitivity and Resistance Testing (DSRT) Oncology Drug Screen

DSRT was performed on PDAC cell lines essentially as we described previously (26,28). Briefly, compounds were dissolved in DMSO and dispensed on tissue culture–treated 384-well plates (Corning) using the Echo 550 (Labcyte Inc.) acoustic liquid handling device. The compounds were plated in five different concentrations in 10-fold dilutions covering a 10,000-fold concentration range appropriate for each compound. Single-cell suspensions (1,000 cells) of each PDAC cell line were transferred to each well using a MultiDrop Combi (Thermo Scientific) peristaltic dispenser, then incubated in a humidified environment at 37°C and 5% CO^2^. After 72 hours, cell viability was measured using CellTiter-Glo luminescent assay (Promega) with a Molecular Devices Paradigm plate reader. Data were normalized to negative control (DMSO only) and positive control wells (100 μmol/L benzethonium chloride, effectively killing all cells). Curve fitting and calculations of drug sensitivity scores were performed as previously described (79). Our analysis was based on the delta drug sensitivity score (dDSS), defined as the difference between the drug sensitivity score (DSS) for the combination treatment and that for ERKi alone (26). A negative score indicates decreased proliferation, with values less than negative five considered synergistic, while a positive score reflects increased proliferation, with values greater than five considered antagonistic.

### CRISPR-Cas9 Loss-of-Function Screen

The CRISPR-Cas9 oncogenic pathway library is comprised of five sgRNAs per gene targeting 2,240 genes encoding components of major cellular and oncogenic signaling networks, 50 control non-targeting sgRNAs and 750 targeting 150 control essential/nonessential genes (29,34). The CRISPR screen was performed as we described recently (80). Briefly, 3.2 × 10^8^ PANC-1 cells were stably infected with lentiviral particles at a multiplicity of infection of 0.2, followed by a growth medium exchange supplemented with 2 µg/ml of puromycin to select for infected cells. After selection, puromycin concentration was halved and cells were cultured for 7 days, allowing for essential genes to be depleted. Cells were then split into 500 cm^2^ tissue culture plates at a concentration of 1.25 × 10^7^ cells per plate and cultured independently throughout the screen. The next day, growth medium supplemented with vehicle (DMSO) or 30 nM RMC-7977 was added to the plates. Cells were maintained for 4 weeks, exchanging RMC-7977/DMSO every 3-4 days and passaging as necessary, maintaining 1.25 × 10^7^ cells per plate (1,000x coverage of each sgRNA) in each technical replicate. After 4 weeks of treatment, cells were harvested for isolation of genomic DNA using the DNeasy Blood and Tissue Kit (Qiagen). The guide RNAs were PCR-amplified, followed by an additional PCR step to add indexing primers. The PCR product was resolved in a 2% agarose gel and purified using the QIAquick gel extraction kit (Qiagen). Deep sequencing was performed with an Illumina NextSeq 500/550 High Output 75 cycle kit on a NextSeq 500 DNA sequencer (Illumina) and enrichment or depletion of sgRNAs was assessed by MAGeCKFlute analysis (81).

### TCGA survival analysis

The association between *AXL* mRNA expression and clinical outcome was evaluated in the TCGA pancreatic adenocarcinoma cohort (TCGA-PAAD) using the Genetic Determinants of Cancer Patient Survival resource (tcga-survival.com; v.2, Smith and Sheltzer, 2022). Pan-Cancer Atlas RNA-sequencing data were combined with clinical outcome data from the TCGA Pan-Cancer Clinical Data Resource, using overall survival as the endpoint. Expression values were log2-transformed, and the association between continuous *AXL* expression and overall survival was assessed by Cox proportional hazards regression. For Kaplan-Meier visualization, patients were dichotomized at the cohort mean expression value and survival distributions were compared by two-sided log-rank test.

### Proliferation Assays

For anchorage-dependent 2D proliferation assays, PDAC cells were seeded in flat bottom 96-well plates (Corning) at a density of 1,000-3,500 cells per well, depending on the cell line. Cells were grown for 24 hours prior to pharmacological treatments. Drugs were dispensed using a D300e Digital Dispenser (Tecan), and the cells were cultured for an additional five days. Following treatment, the cells were either incubated for 10 minutes in medium containing 10 μg/mL DAPI (4’,6’-diamidino-2-phenylindole) and imaged on an Agilent BioTek Cytation 5, or fixed with 70% ethanol fixation at −20°C. Fixed cells were then stained with a 1% propidium iodide (PI) solution in PBS (ThermoFischer, BMS500PI) for 30 minutes at room temperature. Imaging was performed using CQ1 confocal fluorescence microscope (Yokogawa). Cell counts were obtained using either BioTek Gen5 software or CellPathfinder software and normalized to day zero values, which were seeded in a separate plate.

For the spheroid proliferation assay, PDAC cell lines were seeded at a density of 1,500– 6,000 cells per well, depending on the cell line, in 150 µL of the respective growth medium supplemented with 2% growth factor-reduced Matrigel (Corning), and cultured in low-attachment, round-bottom 96-well plates (Corning). The plates were centrifuged at 300 x g for 10 minutes. Cells were grown for 24 hours prior to pharmacological treatments. Drugs were dispensed using the D300e Digital Dispenser (Tecan). After seven days of treatment, cell viability was assessed using the CellTiter-Glo 3D Cell Viability assay (Promega) according to the manufacturer’s recommendation and luminescence was measured with a microplate reader (Synergy). For LUAD cell lines, cells were seeded in ultra-low-attachment 96-well plates (Corning 7007), centrifuged for 10 minutes at 1000 x g, and incubated for 72 hours to form spheroids. Cells were treated with compound or 0.1% DMSO incubated for 5 days. The number of viable cells was assessed via CellTiter-Glo 3D Reagent (Promega) according to manufacturer instructions. For confluency using Incucyte, cells were seeded at 1000 cells per well in tissue-culture treated 96-well plates and the wells were imaged every 4 hours for 4 days on a Sartorious Incucyte Instrument and the confluence of each well was assessed using the accompanying Incucyte software. Data are plotted as the mean ± SEM of 12 replicate wells.

For PDAC organoid proliferation assays, cells were seeded in white-walled, flat-bottom, PolyHEMA-treated 96-well plates using an ice-cold growth medium with 10% Matrigel mixture (Corning). Cell density at seeding was between 2000 and 4000 cells per well, depending on organoid line. Cultures were maintained for 2 days before treatment were added with a D300e Digital Dispenser. After 5 days of treatment, cell viability was assessed with CellTiter-Glo 3D assay (Promega) according to the manufacturer’s recommendation and luminescence was measured with a SpectraMax i3x plate reader.

For colony formation assays, PDAC cells were seeded in 6-well plates (Corning) at a density of 1,000-3,500 cells per well, depending on the cell line. Cells grown for 24 hours prior to pharmacological treatments and allowed to grow for 14 days. Medium was refreshed every 3 days. After 14-days treatment, cells were washed with PBS once and fixed in acetic acid and methanol (1:7 v/v) for 10 minutes at room temperature. Cells were subsequently stained with a crystal violet solution (0.5%) for 2 hours. Images of the dry cell colonies were taken and the area covered was quantified by the software Fiji (ImageJ) using the Plugin ColonyArea. All data were normalized to their corresponding controls.

The growth percentage was calculated by normalizing DMSO control (100% of Growth) for each biological replicate. Treatment response curves were calculated and generated by GraphPad Prism. The standard error of mean was used to generate the error bar between biological replicates. For synergy score analysis, growth analysis data were used to calculate the Bliss synergy score by SynergyFinder (3.16.0) using R (4.6). Heatmaps represent the independent synergy score at each combination.

### Western Blot

Cells were seeded into 6-well plates (Sarstedt, 833.920) at 8 × 10⁴ to 3 × 10⁵ cells per well, depending on treatment duration, and allowed to adhere overnight before the medium was replaced with fresh medium containing the respective treatment. After treatment, cells were washed twice with ice-cold PBS and lysed on ice in Pierce RIPA buffer (Thermo Fisher Scientific, 89900) supplemented with protease and phosphatase inhibitor cocktails (cOmplete EDTA-free, 11836170001; PhosSTOP, 4906837001). Lysates were cleared by centrifugation (13,000 rpm, 10 min, 4°C), and protein concentrations were determined with the Pierce BCA Protein Assay Kit (Thermo Fisher Scientific, 23225) on a Synergy 2 microplate reader (BioTek).

Equal amounts of protein (20–30 µg, depending on the cell line) were mixed with SDS sample buffer containing 100 mM DTT and denatured at 95°C for 5 min. Proteins were resolved on hand-cast 10% polyacrylamide gels (68 V for 40 minutes, then 120 V for 2 hours, on ice) using a Mini-PROTEAN 3 Cell system (Bio-Rad) alongside a PageRuler Prestained Protein Ladder (10–180 kDa), and transferred onto 0.45 µm PVDF membranes (Immobilon-P, Merck, IPVH00010) by wet-tank electroblotting. Membranes were blocked in 5% BSA (Serva, 11926.03) in TBS-T for 1 hour at room temperature and incubated with primary antibodies overnight at 4 °C in 5% BSA/TBS-T. After three washes in TBS-T (10 minutes each), membranes were incubated with HRP-conjugated secondary antibodies in TBS-T for 1 hour at room temperature and washed twice in TBS-T (10 minutes each). Signals were detected with Clarity Western ECL (Bio-Rad, 1705060) or SuperSignal West Femto substrate (Thermo Fisher Scientific, 34094) on a ChemiDoc MP imaging system (Bio-Rad). Band intensities were quantified by densitometry in ImageJ and normalized to β-actin as loading control. The following primary and secondary antibodies were used: anti-actin (rhodamine-conjugated; Bio-Rad #12004164, 1:2500), rabbit anti-phospho-AXL (Cell Signaling Technology [CST] #96453, 1:500), rabbit anti-AXL (CST #8661, 1:2000), rabbit anti-GAS6 (CST #67202, 1:1000), rabbit anti-phospho-ERK1/2 (CST #4370, 1:1000), mouse anti-total ERK1/2 (CST #4696, 1:1000), rabbit anti-phospho-AKT (CST #4060, 1:1000), mouse anti-AKT (CST #2920, 1:1000), rabbit anti-phospho-p90RSK (CST #9344, 1:500), rabbit anti-p90RSK (CST #9355, 1:1000), rabbit anti-MYC (Abcam #ab32072, 1:1000), rabbit anti-phospho-S6 ribosomal protein (CST #4858, 1:1000), mouse anti-S6 ribosomal protein (CST #2317, 1:2000),, rabbit anti-β-actin (CST #4970, 1:2000), anti-mouse IgG (HRP-linked) (CST #7076, 1:2500), and anti-rabbit IgG (HRP-linked) (CST #7074, 1:2500).

H358-GFP or H358-AXL cells were grown in 10 cm dishes, then washed with PBS and lysed with NP-40 Lysis Buffer containing 1x HALT protease/phosphatase inhibitors (Thermo Fisher Scientific). Samples containing 30 µg of protein (assessed by BCA assay, Thermo Fisher Scientific) were resolved on Bis-Tris SDS PAGE gels then transferred to a nitrocellulose membrane and blocked for 1 hour in TBS Intercept Buffer (LiCor). Mouse anti-FLAG (Sigma-Aldrich, F1804) or anti-α-tubulin (Cell Signaling Technology, 3873) at 1:1000 dilution in TBS Intercept Buffer + 0.1% tween-20 was added and incubated at 4°C overnight. After washing a secondary antibody (LiCor, 926-68070) was added at 1:15,000 dilution and incubated for 1 hour at ambient temperature. After washing, the blots were imaged on the LiCor Odyssey.

### Reverse Phase Protein Array

Reverse-phase protein array (RPPA) was done as we have described previously (75). Cell lines (HPAF-II, KP-4, MIA PaCa-2 and Pa16C) were plated onto 6-well plates. The next day cells were treated with vehicle, 10 nM RMC-7977 (RASi), 2 µM bemcentinib (AXLi), or the combination of both for 4, 24, and 72 hours. At the appropriate time point cell lysates were prepared as we described previously (82,83). All treatments were done in biological quadruplicate. The protein concentration was determined using Coomassie Protein Assay Reagent kit (Thermo Fisher Scientific). Protein solution was diluted to 0.5 mg/ml in 2x tris-glycine SDS Sample buffer (Life Technologies) with 5% β-mercaptoethanol and boiled for 8 minutes and stored at −80°C until put on the array. Lysates were immobilized using an Aushon 2470 automated system (Aushon BioSystems) as previously described (82). Samples were printed in three technical replicates together with reference standards for internal controls. To quantify the amount of total protein present in each sample, selected arrays were stained with Sypro Ruby Protein Blot Stain (Molecular Probes) (manufacturing instructions). Remaining arrays were treated with Reblot Antibody Stripping solution (MilliporeSigma) for 15 minutes at room temperature, followed by two washes with PBS, and incubated for 4 hours in I-block (Tropix) prior to immunostaining. Immunostaining was performed using an automated system (Dako Cytomation) previously described. Each array was probed with one antibody targeting the proteins of interest. Arrays were probed with a total of 83 antibodies (Table S1). Each antibody was validated previously for its specificity (84). Biotinylated anti-rabbit (Vector Laboratories, Inc.) or anti-mouse secondary antibodies (DakoCytomation) coupled with a commercially available tyramide-based avidin/biotin amplification system (Catalyzed Signal Amplification System; DakoCytomation) and the IRDye 680RD Streptavidin (LI-COR Biosciences) fluorescence dye were used for the signal amplification and detection. Images for Sypro Ruby and antibody-stained slides were acquired using a Tecan laser scanner (TECAN) and images were analyzed using the commercially available software MicroVigene Version 5.1.0.0 (Vigenetech) (85).

### Animal Models

All mouse experiments were conducted in compliance with ARRIVE guidelines and approved by the Institutional Animal Care and Use Committee (IACUC) of the University of Navarra (CEEA) under protocol R-057–18GN, authorized by the regional Government of Navarra. Mice were randomized for a similar average tumor size across treatment groups at treatment start. Animals were treated without knowledge of anticipated outcomes and blind treatments were followed.

### In Vivo Xenograft Studies

For xenograft experiments, immune-deficient 8- to 12-week-old *Rag2*^−/−^; *ILR2*^−/−^ mice (C;129S4-*Rag2*^tm1.1Flv^ *Il2rg*^tm1.1Flv^/J) from The Jackson Laboratory were used. Human pancreatic cancer cell lines HPAF-II and KP-4 were harvested, washed, and resuspended in 100 μL of phosphate-buffered saline (PBS), then injected subcutaneously into the two lower flanks of mice. Tumor growth was monitored starting one-week post-injection and measured every three days using digital calipers (DIN862, Ref 112-G, SESA Tools) and tumor volume was calculated by the formula: volume = π/6 x length x width^2^. Once tumors reached an average volume of approximately 200 mm³, mice were randomized into treatment groups (*n* = 6 per group). Treatment involved administering bemcentinib at 50 mg/kg (in 0.5% CMC-Na) via oral gavage twice daily, RMC-7977 orally at 3 mg/kg (formulated in 10/20/10/60 (%v/v/v/v) DMSO/PEG400/Solutol HS15/Water) for HPAF-II and 10 mg/kg for KP-4, or a combination of both. Control mice were treated with the corresponding vehicle. All groups were treated for 15 days with tumor volume measurements recorded every 3 days. Subsets of mice (n = 2/group) were sacrificed on treatment day 3 for IHC analyses; remaining mice completed the 15-day course. Following the end of treatment, tumor volume was recorded every 3-5 days, and mice were sacrificed when the tumor size reached 1000 mm³ for HPAF-II and 1400 mm³ for KP-4. Survival endpoint was determined by assessing time-to-event, defined as tumor volume = 800 mm^3^ for HPAF-II and 1000 mm^3^ for KP-4.

All mice were housed in filtered cages of a pathogen-free facility (12 hours light/12 hours dark cycle) and fed with standard diet and water *ad libitum*. All procedures were conducted in compliance with institutional and regulatory guidelines to ensure animal welfare. For the H358-GFP and H358-AXL CDX studies, female athymic nude mice aged 6 weeks were obtained from The Jackson Laboratory (Bar Harbor, ME, USA). All mouse studies and procedures related to animal handling, care, and treatment complied with all applicable regulations and guidelines of the institutional Animal Care and Use Committee (IACUC) of Revolution Medicines with approvals. NCI-H358-GFP and NCI-H358-AXL-OE cells (derived from NCI-H358, ATCC CRL-5807) were cultured as monolayers in RPMI 1640 medium supplemented with 10% fetal bovine serum, 1% penicillin-streptomycin, and 2 µg/mL puromycin. Cultures were maintained at 37°C in a humidified atmosphere containing 5% CO₂. Each mouse was subcutaneously inoculated in the right flank with 1 × 10⁷ cells suspended in 100 µL of a 1:1 mixture of RPMI 1640 and Matrigel. Treatment began upon randomization when tumor reaches an average volume of 149-189 mm³. Tumor-bearing animals were randomized and grouped for treatments (n = 5-6 per group). The vehicle, RMC-7977, and RMC-4998 were administered via daily oral gavage with 10 mL/kg dosing. RMC-7977 and RMC-4998 were formulated in 10/20/10/60 (%v/v/v/v) (DMSO/PEG 400/Solutol HS15/water).

### In Vivo Syngeneic Mouse Studies

Syngeneic subcutaneous experiments were performed in immunocompetent 8-12 week old female F1 C57Bl/6 J x 129S2/Sv mice from Janvier Labs. The murine pancreatic cancer cell line FC1242 (derived from KPC tumors) was harvested and resuspended in PBS. A total of 3×10^5^ cells in 100 μL PBS were injected subcutaneously into both lower flanks. Tumor growth was monitored every 3 days starting at one week post-implantation. When tumors reached ∼200 mm³, mice were randomized into four treatment arms (n = 6 per group) and treated with the same regimens described above: vehicle, bemcentinib (50 mg/kg, twice daily, oral gavage), RMC-7977 (3 mg/kg, daily, oral gavage), or the combination, for 15 days. Following the end of treatment. overall survival was recorded, and mice were sacrificed when the tumor size reached 1000 mm³. Survival endpoint was determined by assessing time-to-event, defined as tumor volume = 800 mm^3^.

Orthotopic tumors were established by injecting 1.5×10^5^ FC1242 cells in 20 μL PBS into the pancreas of 8- to 12-week-old female F1 C57Bl/6 J x 129S2/Sv mice. Tumor burden was monitored weekly using micro computed tomography (microCT). When tumors reached 250-300 mm³ (survival studies) or 100-120 mm³ (immunophenotyping), mice in each independent experiment were randomized into four treatment groups (*n* = 6 per group) and treated with vehicle, bemcentinib (50 mg/kg, twice daily, oral gavage), RMC-7977 (3 mg/kg, daily, oral gavage), or the combination. For survival studies, mice were treated for 15 days and overall survival was recorded. For immunophenotyping, mice were treated for 10 days prior to tumor collection.

### Immunohistochemistry

Immunohistochemical staining was performed using the EnVision TM + System (Dako, K400311-2; Glostrup, Denmark) according to the manufacturer’s recommendations. Paraffin sections (3 µm thick) were cut, dewaxed and hydrated. Antigen retrieval was performed for 30 minutes at 95°C in 0.01 M Tris-1 mM EDTA buffer (pH 9) in a Pascal pressure chamber (S2800, Dako). Slides were allowed to cool for 20 minutes, then endogenous peroxidase was blocked with 3% H_2_ O_2_ in deionized water for 12 minutes and sections were washed in TBS-0.05% Tween 20 (TBS-T). Sections were incubated overnight at 4°C with primary antibodies: anti-phospho-ERK1/2 (Cell Signaling, #4376; 1:400), anti-cleaved caspase-3 (Cell Signaling, #9661; 1:100), anti-Ki67 (Abcam, #ab16667; 1:100), Phospho-histone H3 (Ser10) (Cell Signaling, #9701; 1:200) and anti-cMyc (Abcam, #ab32072; 1:200). After rinsing in TBS-T, sections were incubated with goat anti-rabbit labelled polymer for 30 minutes at room temperature and peroxidase activity was revealed using DAB+ (Dako, #K346811-2). Finally, sections were lightly counterstained with Harris hematoxylin, dehydrated, and coverslipped. Tissue slides were scanned at ×10 magnification and digitized using the Aperio Scan-Scope XT Slide Scanner (Aperio Technologies).

H358-GFP and H358-AXL CDX tumor fragments were fixed in 10% neutral buffered formalin at room temperature for up to 24 hours, stored at 70% ethanol and then processed into paraffin blocks. FFPE sections (4 μm) were stained on the BOND-III Automated IHC Stainer using the manufacturer’s recommended settings. Antigen retrieval was performed using pH 6.2 Epitope Retrieval solution. AXL rabbit monoclonal antibody (Cell Signaling Technology, #8661) with pH 6.2 HIER was used at a dilution of 1:600. Detection was carried out using Biocare MACH4 HRP polymer to detect the primary antibodies. DAB-stained slides were scanned and digitized with a 3DHISTECH Panoramic MIDI whole-slide scanner at 1.5× magnification.

### RNA Extraction

Tumor samples were excised from xenografted mice, immediately snap-frozen in liquid nitrogen, and stored at −80°C until processing. For RNA extraction, ∼50 mg of tumor tissue was transferred to a pre-chilled microcentrifuge tube containing 800 μL of TRIzol reagent (Thermo Fisher Scientific). The tissue was mechanically disrupted and homogenized using a sterile pestle until fully lysed, ensuring complete tissue dissociation. The homogenate was incubated at room temperature for 5 minutes to allow nucleoprotein complexes to dissociate. Following this, 200 μL of chloroform was added, and the mixture was vortexed for 15 seconds and incubated at room temperature for 5 minutes. The sample was centrifuged at 12,000 × g for 15 minutes at 4°C to separate phases. The aqueous phase containing RNA was carefully transferred to a fresh tube, and RNA was precipitated by adding 0.5 mL of isopropanol. The RNA pellet was washed twice with 75% ethanol, air-dried, and resuspended in RNase-free water. The RNA was then further purified using the RNeasy kit (Qiagen ID: 74104) according to the manufacturer’s instructions, to ensure high purity and removal of contaminants. RNA quality and concentration were evaluated using a NanoDrop spectrophotometer. Purified RNA was stored at −80°C for RNA sequencing

### CDX RNA Sequencing

RNA was subjected to quantity and quality control using Qubit HS RNA Assay Kit (Thermo Fisher Scientific) and 4200 Tapestation with High Sensitivity RNA ScreenTape (Agilent Technologies). All RNA samples were high-quality, with RIN values higher than 7. Library preparation was performed using the Illumina Stranded mRNA Prep Ligation kit (Illumina) following the manufacturer’s protocol. All sequencing libraries were constructed from 100 ng of total RNA according to the manufacturer’s instruction. Briefly, poly(A) containing RNA molecules were selected and purified using magnetic beads coated with poly(T) oligos. Poly(A)-RNAs were fragmented and reverse transcribed into first cDNA strand using random primers. The second cDNA strand was synthesized in the presence of dUTP to ensure strand specificity. Resulting cDNA fragments were purified with AMPure XP beads (Beckman Coulter), adenylated at 3′ ends and then ligated with uniquely indexed sequencing adapters. Ligated fragments were purified and PCR amplified to obtain the final libraries. The quality and quantity of the libraries were verified using Qubit dsDNA HS Assay Kit (Thermo Fisher Scientific) and 4200 Tapestation with High Sensitivity D1000 ScreenTape (Agilent Technologies). Libraries were then sequenced using a NextSeq2000 sequencer (Illumina). 20-30 million pair-end reads (100 bp; Rd1:51; Rd2:51) were sequenced for each sample and demultiplexed using bcl2fastq.

### RNA-seq Data Analyses

TrimGalore (v0.6.6) was used for QC and trimming reads with a quality threshold (phred33) of 28, retaining paired reads with at least 35 high quality basepairs in each read. Transcriptomes were aligned using the STAR (v2.6.1c) (86) alignment tool with Gencode3 reference GRCh38.p13 and annotation set v43 (87). Transcripts were quantified and summarized per gene using Salmon (v0.11.3) (88) followed by tximport (v1.32.0). Ensembl annotations were added for each gene using biomaRt (v2.60.1) (89,90). Transcripts of tumor models were independently filtered for genes where at least 20 reads were detected in at least four samples, removing read-through transcripts and non-protein coding transcripts, and independently normalized using the TMMwsp method in EdgeR (v4.2.2) (91). Differential gene expression statistics were calculated with the glmQLFTest function in EdgeR. GSEA was computed with fGSEA (v1.30.0) for Hallmark (60), Klomp_KRAS_ERK10 (9) and Hibshman_MYC (92) gene sets. Gene essentiality was approximated as beta values from DepMap (24Q4)12 CRISPR *KRAS*-mutant PDAC cell line analyses and used to compare differentially expressed genes between HPAF-II and KP-4 tumor models.

### Tumor Immunophenotyping by Multiparametric Flow Cytometry

Single-cell suspensions from FC1242 orthotopic tumors were prepared by enzymatic digestion with 1 mg/mL collagenase D (Sigma-Aldrich, #11088858001) and 50 μg/mL DNase I (Roche, #11284932001) at 37°C for 30 minutes, followed by mechanical dissociation through 70 μm cell strainers. Erythrocytes were lysed using RBC lysis buffer (Gibco) and leukocytes were enriched over 35% Percoll (GE Healthcare). Dead cells were excluded using PromoFluor-840 NIR Maleimide (PromoCell). Fc receptors were blocked with anti-mouse CD16/CD32 (BD Biosciences, #553142, 1:200). Cells were stained with lymphoid and myeloid antibody panels.

The lymphoid panel included: CD45-APC-H7 (BioLegend, #103116, 1:500), CD3e-BV395 (BD Biosciences, #563565, 1:80), CD8-BUV395 (BD Biosciences, #563786, 1:200), CD4-BUV496 (BD Biosciences, #612952, 1:200), CD44-BV510 (BioLegend, #103044, 1:200), CD62L-AF700 (BioLegend, #104426, 1:800), CD25-PerCP/Cy5.5 (BioLegend, #102028, 1:50), CD19-BUV661 (BD Biosciences, #612971, 1:200), NKp46-APC (BioLegend, #137608, 1:20), LAG3-BV650 (BioLegend, #125227, 1:400), GITR-FITC (BioLegend, #126308, 1:100), and PD-1-BV421 (BioLegend, #135221, 1:20).

The myeloid panel included: CD11b-BUV661 (BD Biosciences, #612977, 1:160), CD11c-PE (BD Biosciences, #557401, 1:100), Ly6G-FITC (BioLegend, #127605, 1:200), Ly6C-BV510 (BioLegend, #128033, 1:200), F4/80-APC (BioLegend, #123116, 1:80), MHC-II-BV650 (BioLegend, #107641, 1:80), CD38-PerCP/Cy5.5 (BioLegend, #102722, 1:160), and CD206-PE/Cy7 (BioLegend, #141720, 1:40).

For intracellular targets, cells were fixed and permeabilized using the eBioscience Fixation/Permeabilization Kit (Thermo Fisher Scientific) according to the manufacturer’s instructions and stained intracellularly with FoxP3-PE/Cy7 (Abcam, #ab210232, 1:100) for the lymphoid panel and CD206-PE/Cy7 (BioLegend, #141720, 1:40) for the myeloid panel. Cells were acquired on a CytoFLEX LX flow cytometer (Beckman Coulter) and data were analyzed using CytExpert software (RRID: SCR_017217). Events were normalized to tumor mass and expressed as events per gram of tumor tissue.

## Data availability

The bulk RNA-sequencing data generated in this study have been deposited at NCBI - SRA with accession number PRJNA1481172.

## Authors’ Contributions

**Y.M. Ching:** Conceptualization, formal analysis, data curation, investigation, writing-review and editing. **S. Narayanan:** Formal analysis, data curation, investigation, writing-review and editing. **J.A. Klomp:** Formal analysis and investigation, writing-review and editing. **T. Isermann:** Formal analysis and investigation. **S. Loewe:** Formal analysis and investigation. **W.-H. Chang:** Formal analysis and investigation. **A.M. Waters:** Formal analysis and investigation. **S.R. Nicewarner Peña:** Formal analysis, investigation. **E. Baldelli:** Formal analysis, investigation. **A.C. Edwards:** Formal analysis and investigation. **T. Boerding:** Formal analysis and investigation. **R. Yang:** Formal analysis and investigation. **C.M. Goodwin:** Formal analysis and investigation. **P. Gautam:** Formal analysis and investigation. **M. Ponz-Sarviśe:** Formal analysis and investigation. **D. Horst:** Formal analysis and investigation. **K. Seamon:** Formal analysis and investigation. **Y. Zhuang:** Formal analysis and investigation. **L. Tran:** Formal analysis and investigation. **J. Jiang:** Conceptualization and supervision. **M. Singh:** Conceptualization and supervision. **K. Wennerberg:** Conceptualization and supervision. **E.F. Petricoin III:** Resources, supervision, and funding acquisition. **K.L. Bryant:** Supervision and funding acquisition. **C.A. Stalnecker:** Formal analysis and investigation. **H.S. Earp:** Supervision and funding acquisition. **A.D. Cox:** Conceptualization, supervision, funding acquisition, writing-review and editing. **C. Sers:** Conceptualization, supervision, funding acquisition, writing-review and editing. **S. Vicent:** Conceptualization, supervision, funding acquisition, writing-review and editing. **B. Papke:** Conceptualization, formal analysis, resources, supervision, funding acquisition, writing-review and editing. **C.J. Der:** Conceptualization, formal analysis, resources, supervision, funding acquisition, writing-review and editing.

## Acknowledgments

We thank Kris Wood (Duke) for the druggable genome CRISPR library, Dr. Anirban Maitra (MD Anderson Cancer Center) for providing PDAC cell lines, Drs. David Tuveson (Cold Spring Harbor Laboratory) and Calvin Kuo (Stanford) for PDAC organoids. W.H. Chang was supported by the American Cancer Society (ACS) PF-23-1072348-01-CDP. A.M. Waters was supported by the National Cancer Institute (NCI) K22CA276632, ACS IRG-23-1141524, Pancreatic Cancer Action Network grant 26-RRG-WATE, and the Gregory Lawton Pancreatic Cancer Research Fund, and Project Purple and Ohio Cancer Research grants. K. Wennerberg was supported by NCI P01CA203657 and P50CA257911. E.F. Petricoin was supported by P01CA203657. K.L. Bryant was supported by NCI P50CA257911 and R37CA251877 and Department of Defense W81XWH2110693. C.A. Stalnecker was supported by NCI T32CA009156 and F32CA232529. H.S. Earp received support from NCI R01CA270792 and P50CA257911. S. Vicent is supported by a Project PID2023-152755OB-I00 funded by MICIU/AEI/10.13039/501100011033 and by FEDER, UE. Support was provided by NCI P50CA257911 and R35CA232113 to A.D. Cox and/or C.J. Der, to C.J. Der by Pancreatic Cancer Action Network (22-WG-DERB), and the Department of Defense W81XWH2110692. C.J. Der and C. Sers were supported from the Einstein Foundation (EVF-2018-431-2) and Deutsche Krebshilfe 70117399. C. Sers received funding from Deutsche Krebshilfe 70114756 and Deutsche Forschungsgemeinschaft (DFG) FOR5628 (513752256). B. Papke was supported by the Deutsches Konsortium für Translationale Krebsforschung (DKTK Berlin) (BE01-SOB-FON-PAP and BE01-SOB-YAC-PAP).

The authors used ChatGPT and Claude for language editing of author-written text, including wording, grammar, spelling, punctuation, and readability. The authors reviewed and edited all AI-assisted text and take full responsibility for the final content of the manuscript.

