## Supplementary figures for "Concurrent AXL Inhibition Enhances RAS and ERK Inhibitor Efficacy in *KRAS*-mutant Pancreatic and Lung Cancer"

**Supplementary Figure S1**

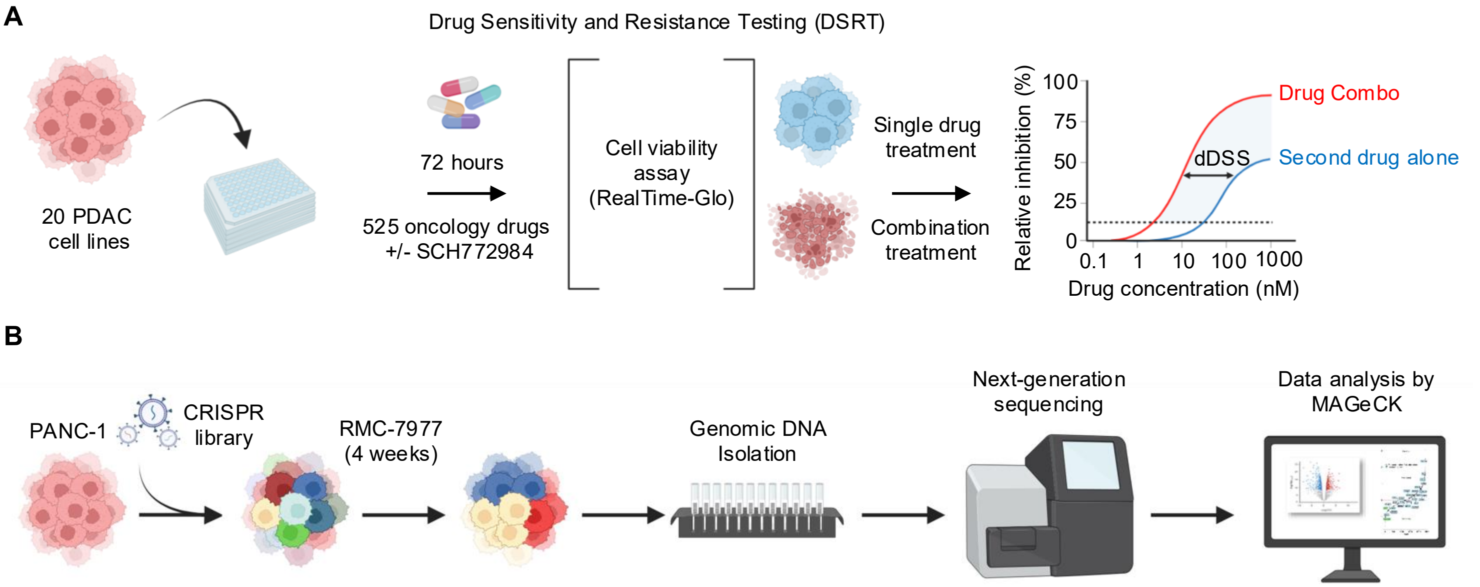


**Supplementary Figure S1.** Unbiased chemical and genetic functional screens to identify modulators of sensitivity to RAS-ERK inhibition. **A,** Schematic overview of the Drug Sensitivity and Resistance Testing (DSRT) workflow. **B,** Schematic overview of the CRISPR-Cas9 loss of function screen workflow with druggable genome library targeting 2,240 genes. Lentiviral particles carrying the library were transduced into PANC-1 cells, followed by puromycin selection and gene editing via Cas9 with sgRNAs for seven days. Cells were then cultured and treated with either DMSO (control) or RASi (RMC-7977, 30 nM). After four weeks, genomic DNA (gDNA) was harvested, sequenced, and analyzed.

**Supplementary Figure S2**

**A**

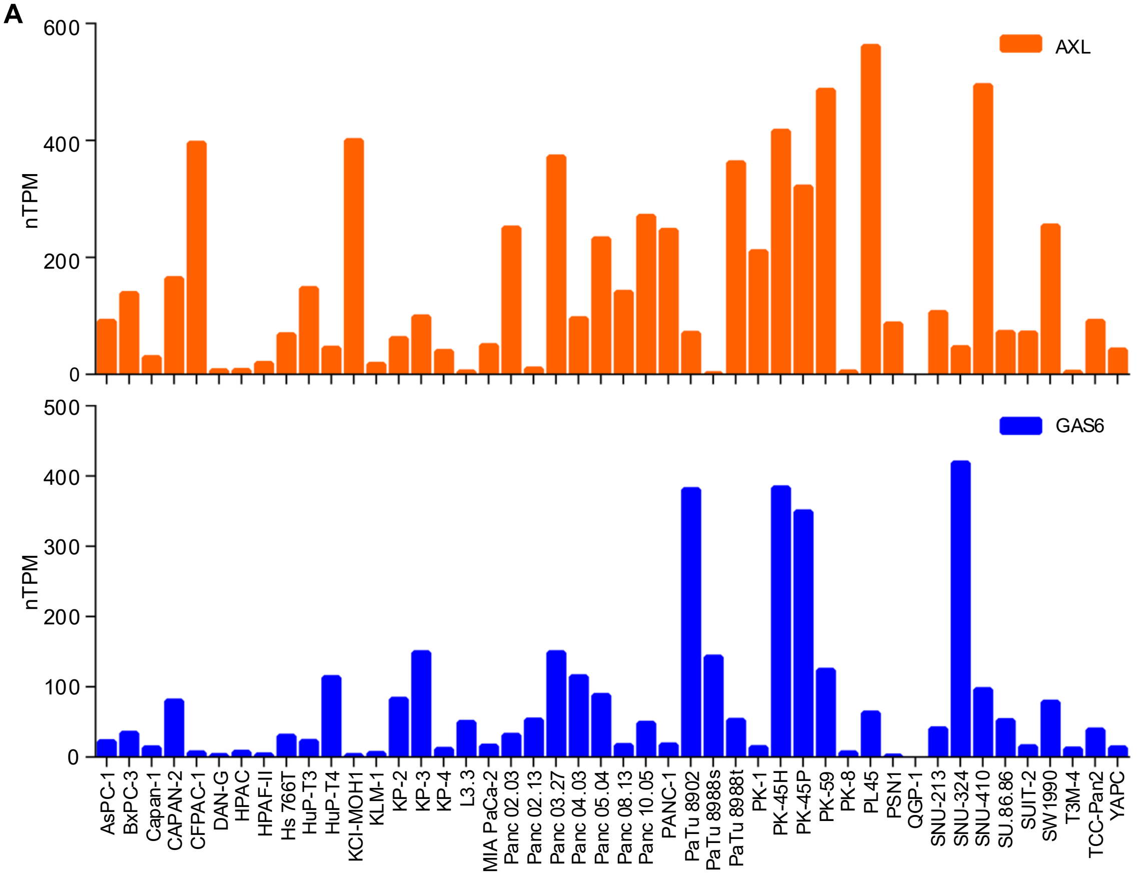


**B**

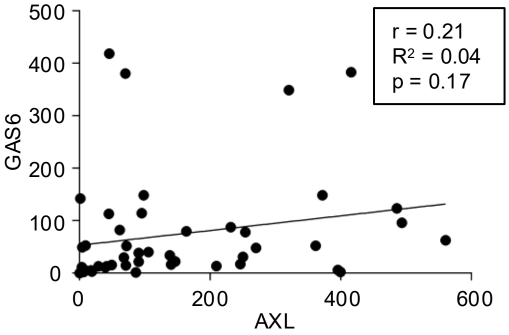


**C**

| Cell line | $GI_{50}$ (nM) |
|---|---|
| BxPC-3 | 601 |
| CFPAC | 487 |
| HPAC | 414 |
| HPAF-II | 462 |
| KP-4 | 394 |
| MIA PaCa-2 | 1036 |
| Pa01C | 346 |
| Pa14C | 686 |
| Pa16C | 706 |
| PANC-1 | 1386 |
| SW1990 | 260 |
| UM53 | 732 |

**D**

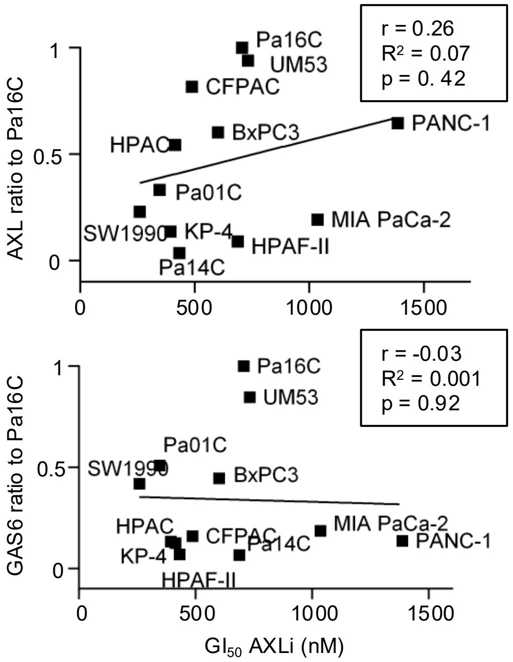


**E**

| Cell line | Compound | $GI_{50}$ (nM) |
|---|---|---|
| PANC-1 | UNC2025 | 141 |
| | Bemcentinib | 959 |
| SW1990 | UNC2025 | 117 |
| | Bemcentinib | 512 |
| Pa14C | UNC2025 | 154 |
| | Bemcentinib | 532 |

**Supplementary Figure S2**

**F**

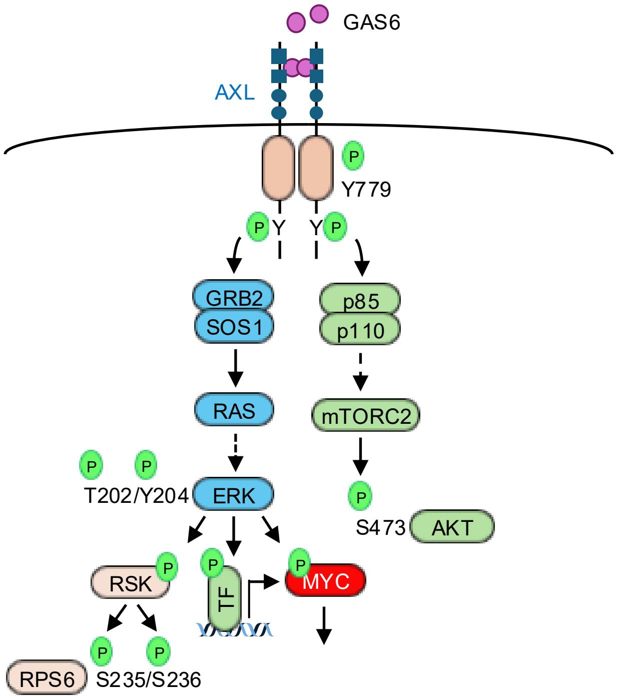


**Supplementary Figure S2.** Correlation Analysis of AXL and GAS6 Expression with AXLi Sensitivity and ERKi Response in PDAC. **A,** RNA analysis of *AXL* (top) and *GAS6* (bottom) normalized transcripts per million (nTPM) in a panel of 46 PDAC cell lines (data from protein atlas). **B,** Pearson correlation analysis of *GAS6* versus *AXL* RNA level from **A** (r = 0, no correlation; r = ±1, positive or negative correlation) **C,** $GI_{50}$ values from proliferation assay, 5 days, $GI_{50} < 600$ nM (sensitive), 600 nM< $GI_{50} < 750$ nM (moderate) and > 750 nM (resistance) to AXLi, measured at Charité Universitätsmedizin Berlin. **D,** Pearson correlation analysis from AXL (top) or GAS6 (bottom) expression level versus $GI_{50}$ values of AXLi in indicated PDAC cell lines. (r = 0, no correlation; r = ±1, positive or negative correlation) **E,** Comparison of the AXLi and UNC2025 in PDAC cell lines after 5 days measured at the University of North Carolina at Chapel Hill. **F,** GAS6-AXL signaling schematic with its two main downstream effector pathways.

**Supplementary Figure S3**

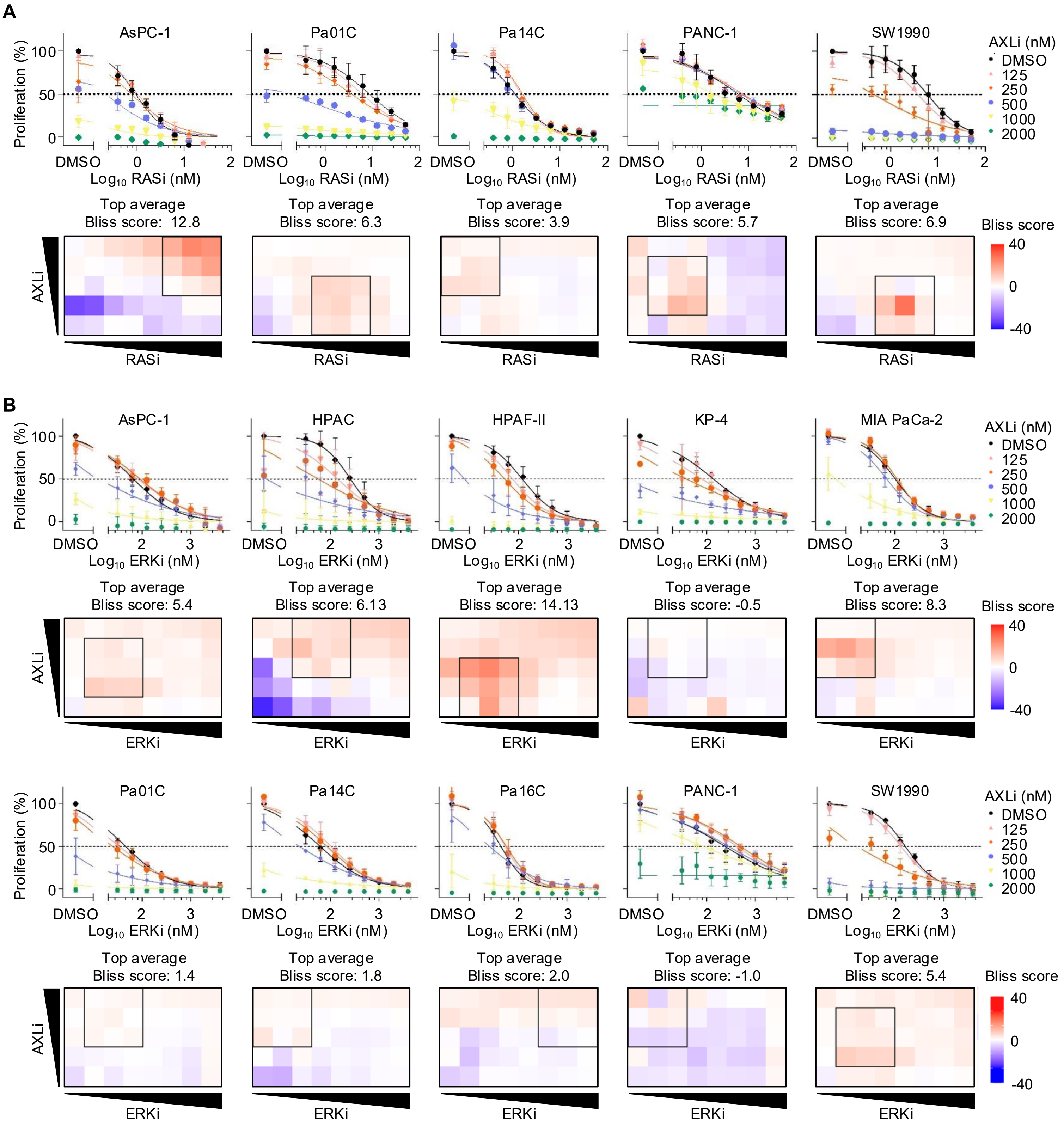

**Supplementary Figure S3**

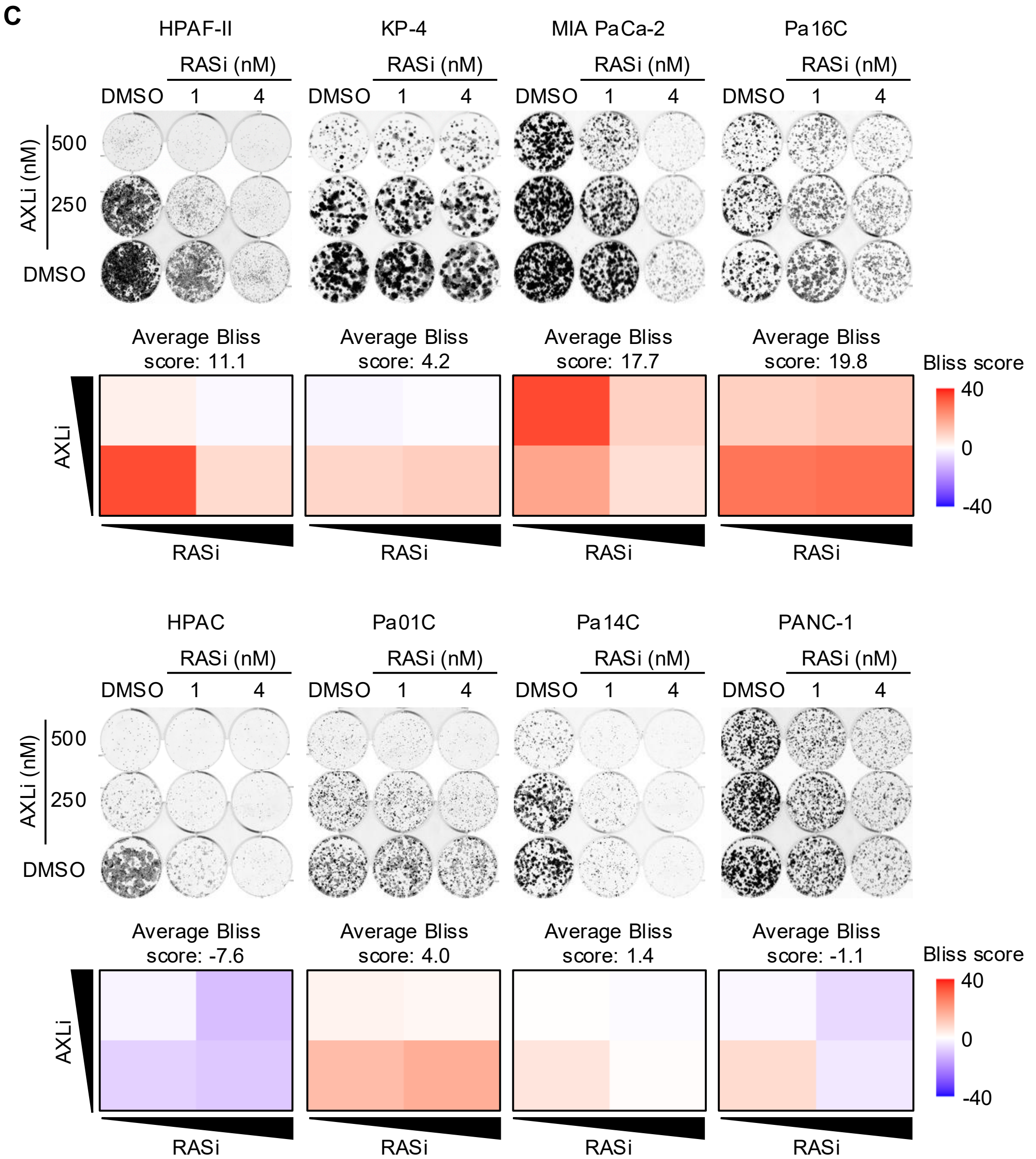


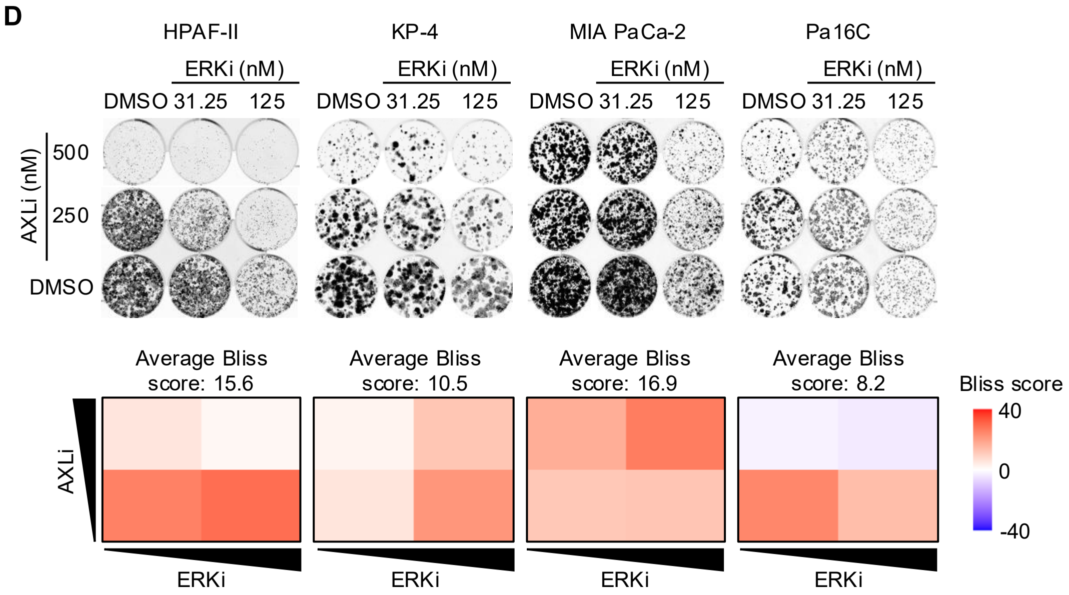

### Supplementary Figure S3

**D (cont.)**

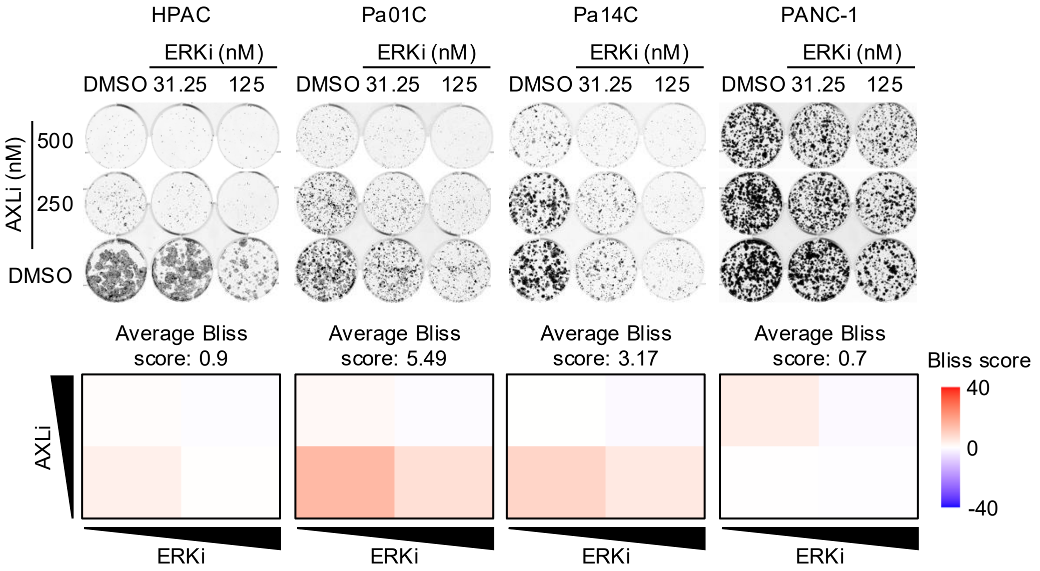


**E**

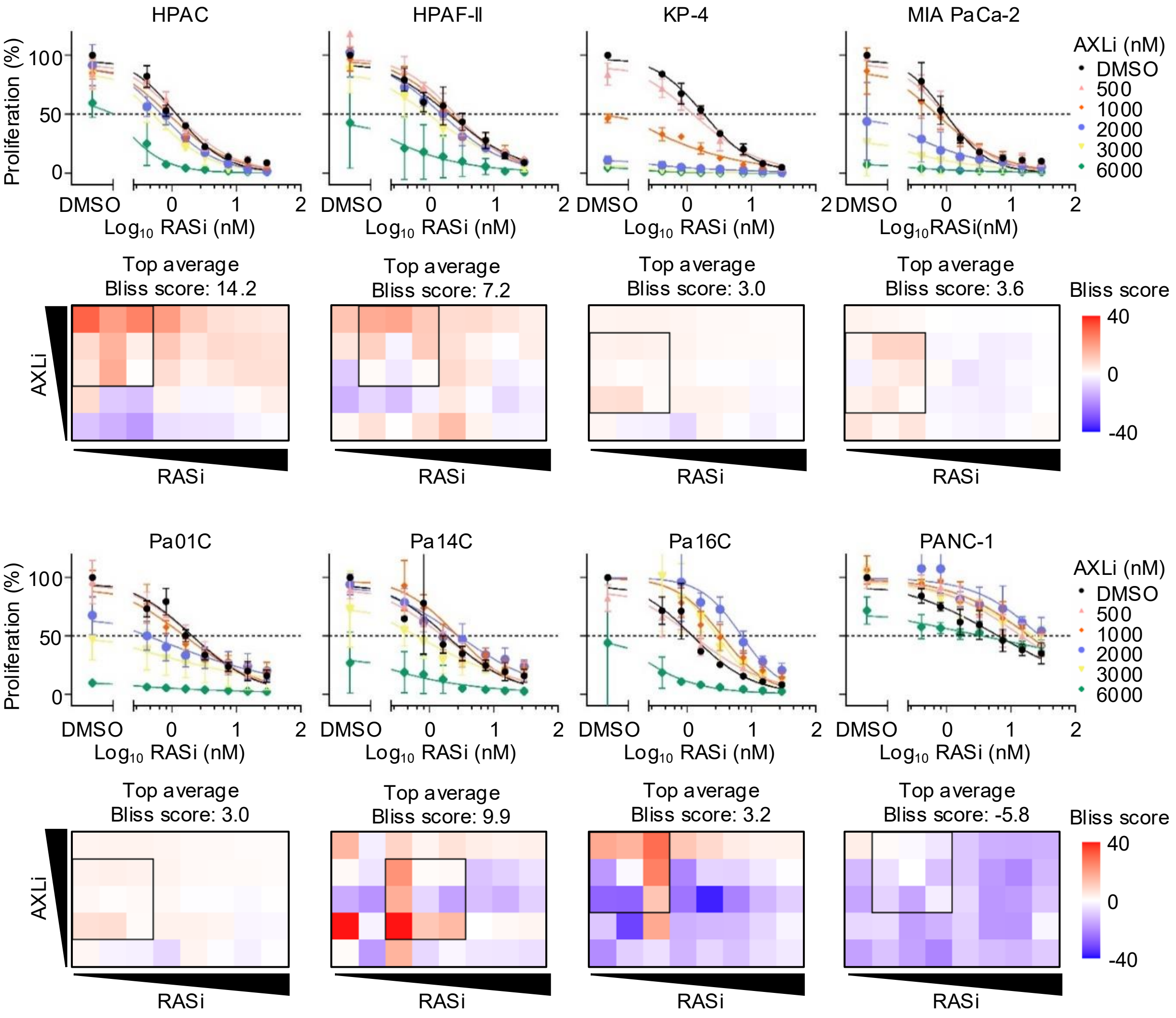

**Supplementary Figure S3**

**F**

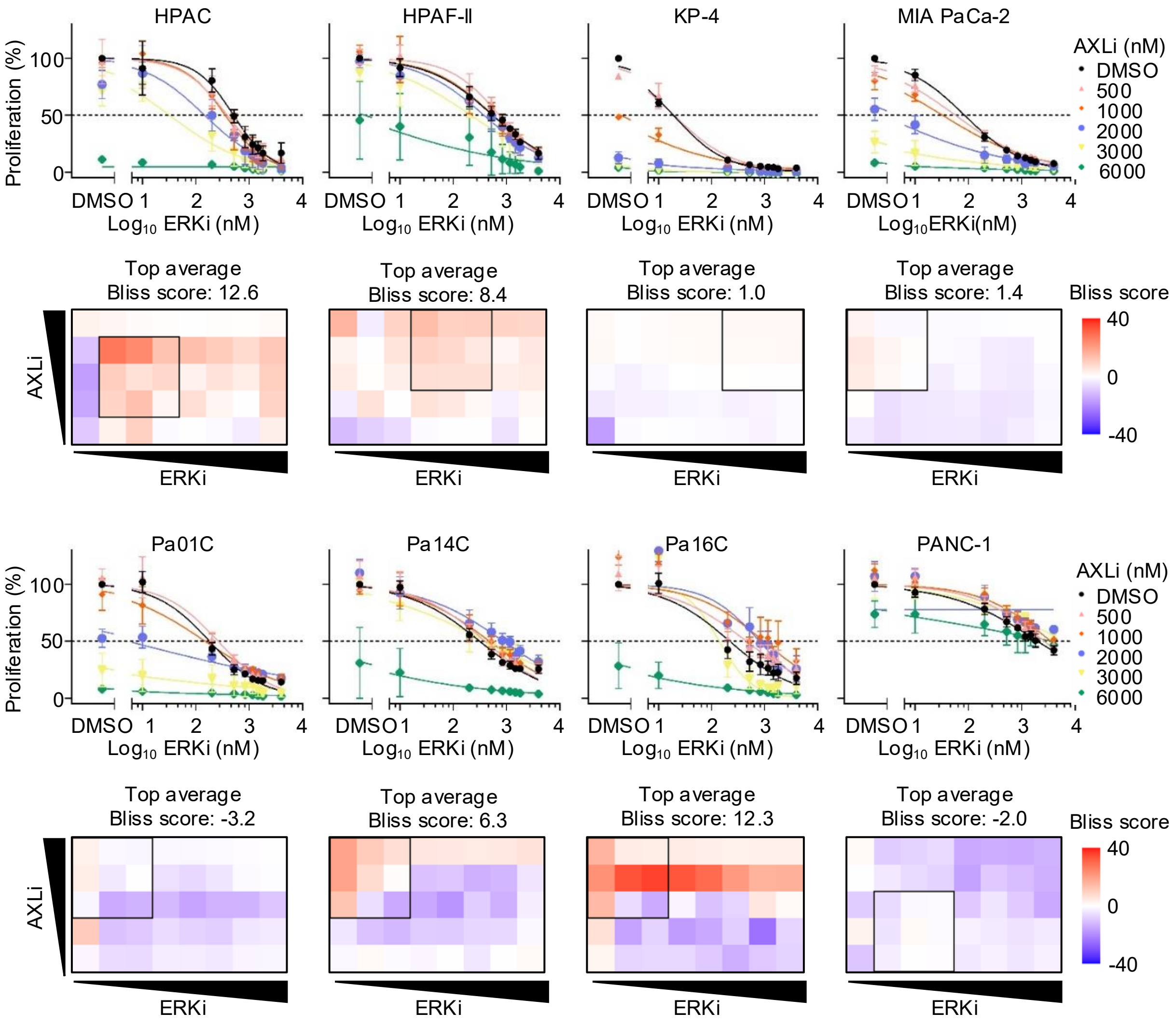


**G**

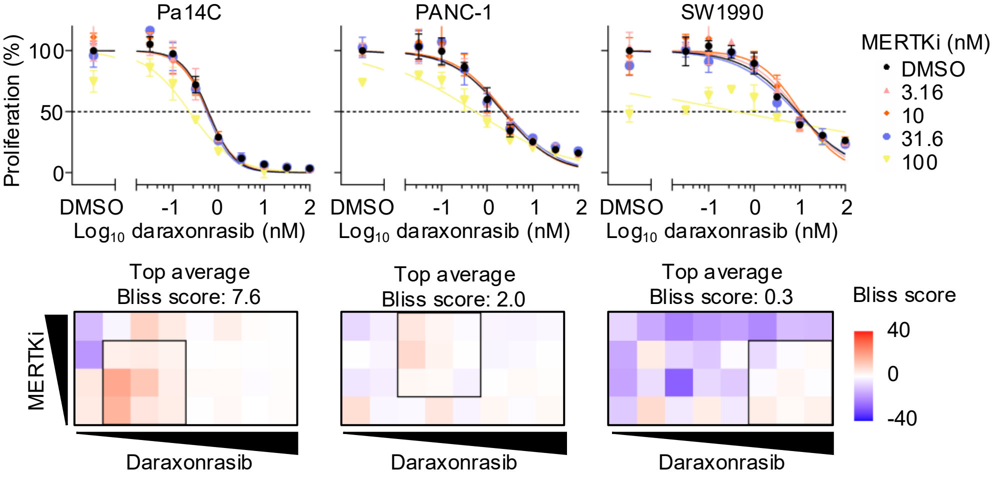

### Supplementary Figure S3

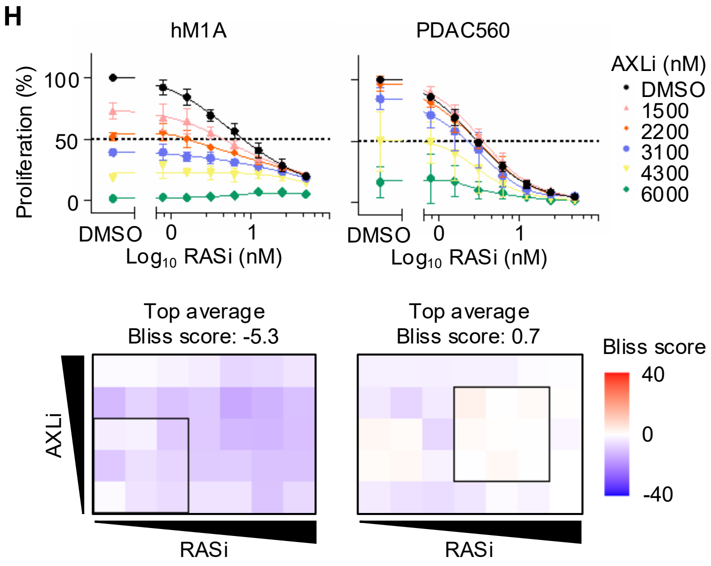


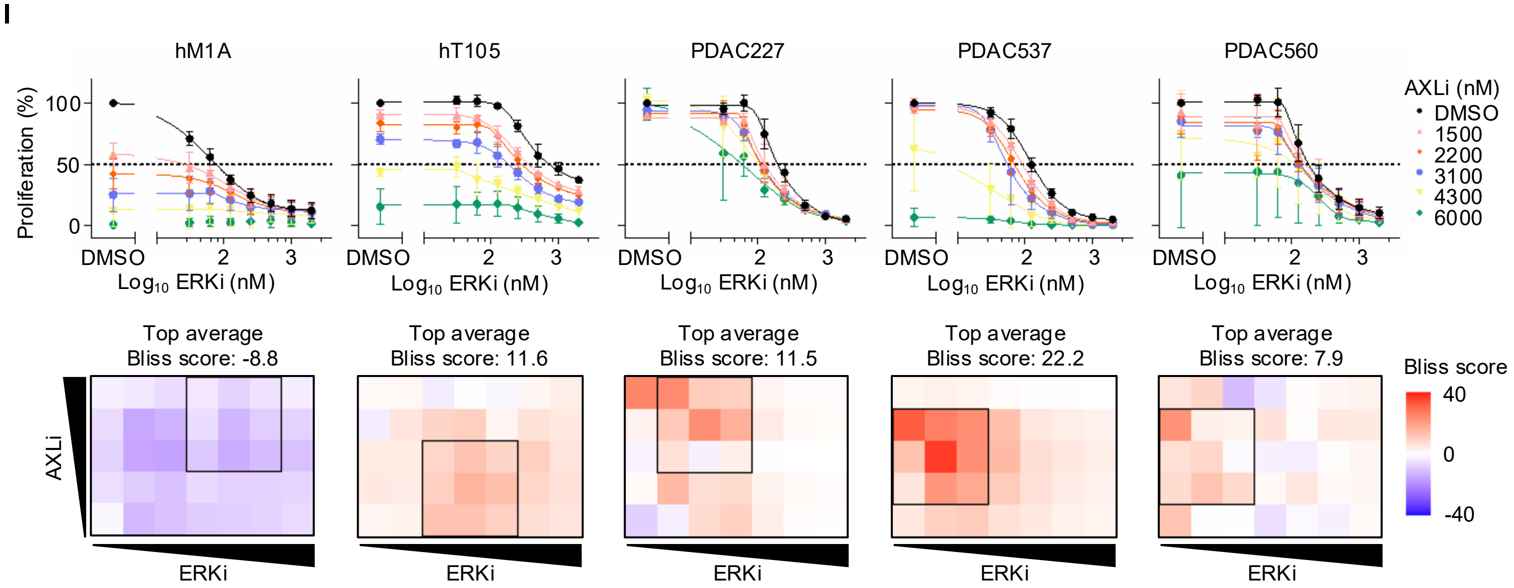

**Supplementary Figure S3**

J

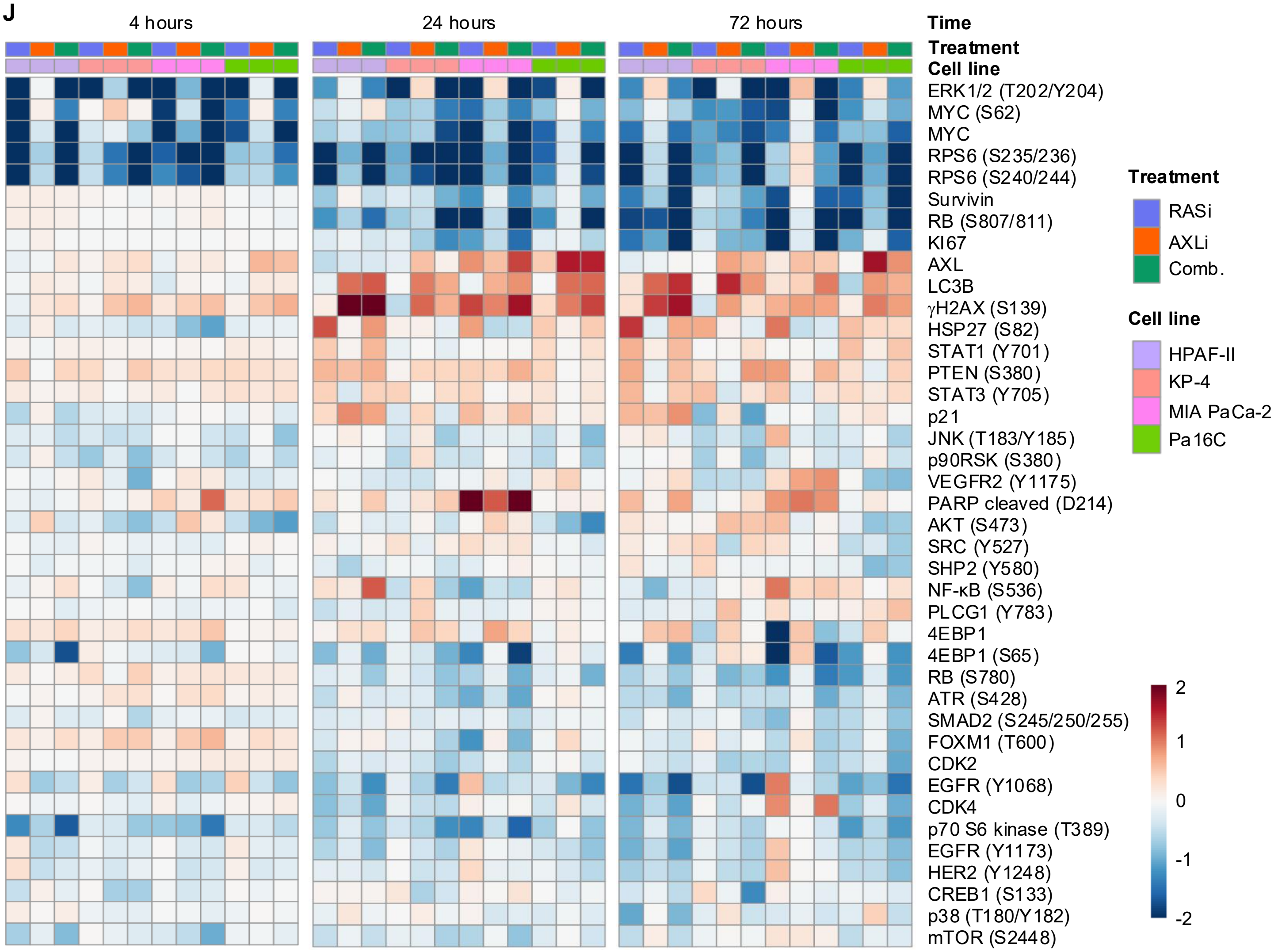


K

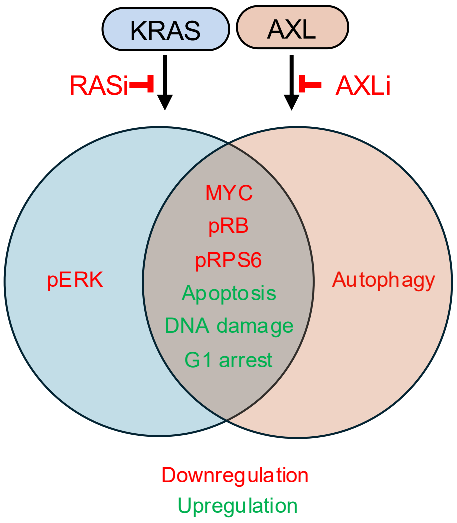

**Supplementary Figure S3.** Combined AXLi and RASi / ERKi treatment synergistically inhibit PDAC growth *in vitro*. **A,** Upper panels: Proliferation of PDAC cell lines treated with increasing concentrations of RASi and AXLi (125-2000 nM) for 5 days, shown as percent proliferation relative to DMSO; the dashed line indicates $GI_{50}$. Lower panels: BLISS synergy heatmaps (red, synergy; blue, antagonism; scale −40 to +40), with the top-average BLISS score and the highest-synergy 3×3 region (black outline) indicated ($n$ = 3 biological replicates). **B,** Proliferation of PDAC cell lines treated with increasing concentrations of ERKi and AXLi (125-2000 nM) for 5 days, shown as percent proliferation relative to DMSO; the dashed line indicates $GI_{50}$. Lower panels: BLISS synergy heatmaps (red, synergy; blue, antagonism; scale −40 to +40), with the top-average BLISS score and the highest-synergy 3×3 region (black outline) indicated ($n$ = 3 biological replicates). **C,** Long-term clonogenic colony formation assays with PDAC lines treated with increasing concentrations of RASi (1 and 4 nM) and AXLi (250 and 500 nM) for 14 days ($n$ = 3 biological replicates), Lower panels: BLISS synergy heatmaps based on BLISS synergy scores calculated from colony area presented above in upper panel using synergyfinder (red, synergy; blue, antagonism; scale −40 to +40), **D,** Long-term colony formation assays treated with increasing concentrations of ERKi (31.25 and 125 nM) and AXLi (250 and 500 nM) for 14 days ($n$ = 3 biological replicates), Lower panels: BLISS synergy heatmaps based on BLISS synergy scores calculated from colony area presented above in upper panel using synergyfinder (red, synergy; blue, antagonism; scale −40 to +40), **E,** Proliferation of the indicated PDAC cell line 3D spheroid cultures treated with increasing concentrations of RASi and AXLi (500-6000 nM) for 7 days, shown as percent proliferation relative to DMSO; the dashed line indicates $GI_{50}$. Lower panels: BLISS synergy heatmaps (red, synergy; blue, antagonism; scale −40 to +40), with the top-average BLISS score and the highest-synergy 3×3 region (black outline) indicated ($n$ = 3 biological replicates). **F,** Proliferation of the indicated PDAC cell line 3D spheroid cultures treated with increasing concentrations of ERKi and AXLi (500-6000 nM) for 7 days, shown as percent proliferation relative to DMSO; the dashed line indicates $GI_{50}$. Lower panels: BLISS synergy heatmaps (red, synergy; blue, antagonism; scale −40 to +40), with the top-average BLISS score and the highest-synergy 3×3 region (black outline) indicated ($n$ = 3 biological replicates). **G,** Upper panels: Proliferation of the indicated PDAC cell lines (Pa14C, PANC-1, SW1990), treated with increasing concentrations of daraxonrasib and MERTKi (3.16 nM-100 nM) for 5 days, shown as percent proliferation relative to DMSO; the dashed line indicates $GI_{50}$. Lower panels: BLISS synergy heatmaps (red, synergy; blue, antagonism; scale −40 to +40), with the top-average BLISS score and the highest-synergy 3×3 region (black outline) indicated ($n$ = 3 biological replicates). **H,** Upper panels: Proliferation of PDAC 3D organoid models (hM1A, PDAC560) treated with increasing concentrations of RASi and AXLi (1500-6000 nM) for 5 days, shown as percent proliferation relative to DMSO; the dashed line indicates $GI_{50}$ ($n$ = 3 biological replicates). Lower panels: BLISS synergy heatmaps (red, synergy; blue, antagonism; scale −40 to +40), with the top-average BLISS score and the highest-synergy 3×3 region (black outline) indicated. **I,** Upper panels: Proliferation of PDAC 3D organoid cell culture models treated with increasing concentrations of ERKi and AXLi (1500-6000 nM) for 5 days, shown as percent proliferation relative to DMSO; the dashed line indicates $GI_{50}$ ($n$ = 3 biological replicates). Lower panels: BLISS synergy heatmaps (red, synergy; blue, antagonism; scale −40 to +40), with the top-average BLISS score and the highest-synergy 3×3 region (black outline) indicated. **J,** RPPA analysis of acute (4 hours), intermediate (24 hours), and prolonged (72 hours) response to AXLi (2 μM), RASi (10 nM), and the combination in PDAC cell lines. **K,** Venn diagram of suppressed genes and signaling pathways of RASi, AXLi and the combination.

### Supplementary Figure S4

A

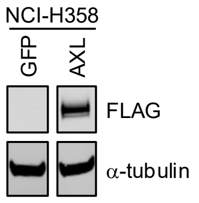


B

| Drug | $GI_{50}$ (nM) | | Resistance |
|---|---|---|---|
| | H358-AXL | H358-GFP | AXL/GFP |
| Adagrasib | 852.5 | 5.2 | 164 |
| RMC-4998 | 304.8 | 0.36 | 847 |
| RASi | 14.8 | 0.48 | 31 |

C

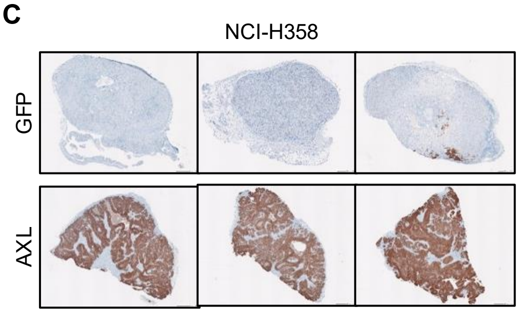


D

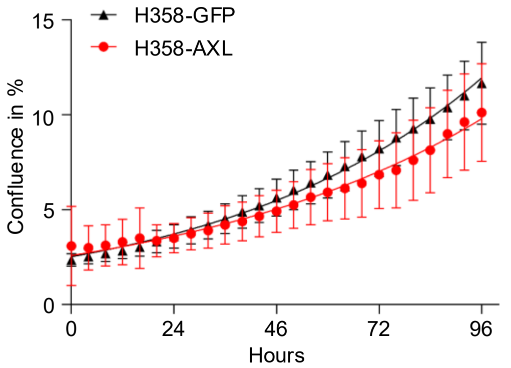


E

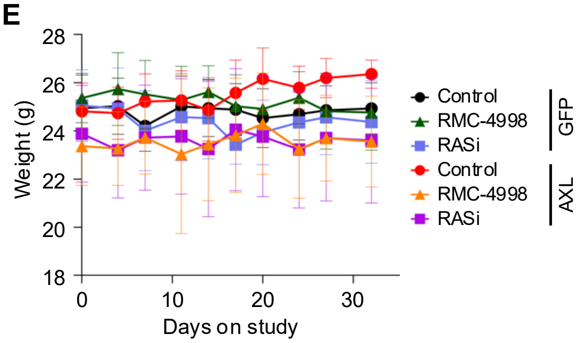


F

| Tumor volume change ratio | |
|---|---|
| Comparison | Significance |
| H358-AXL (RMC-4998) vs. H358-AXL (RASi) | n.s. |
| H358-AXL (RMC-4998) vs. H358-AXL (Vehicle) | **** |
| H358-AXL (RMC-4998) vs. H358-GFP (RMC-4998) | ** |
| H358-AXL (RMC-4998) vs. H358-GFP (RASi) | ** |
| H358-AXL (RMC-4998) vs. H358-GFP (Vehicle) | n.s. |
| H358-AXL (RASi) vs. H358-AXL (Vehicle) | **** |
| H358-AXL (RASi) vs. H358-GFP (RMC-4998) | n.s. |
| H358-AXL (RASi) vs. H358-GFP (RASi) | n.s. |
| H358-AXL (RASi) vs. H358-GFP (Vehicle) | **** |
| H358-AXL (Vehicle) vs. H358-GFP (RMC-4998) | **** |
| H358-AXL (Vehicle) vs. H358-GFP (RASi) | **** |
| H358-AXL (Vehicle) vs. H358-GFP (Vehicle) | **** |
| H358-GFP (RMC-4998) vs. H358-GFP (RASi) | n.s. |
| H358-GFP (RMC-4998) vs. H358-GFP (Vehicle) | **** |
| H358-GFP (RASi) vs. H358-GFP (Vehicle) | **** |

**Supplementary Figure S4.** AXL overexpression confers resistance to RAS inhibition and creates a synergistic dependency on combined AXL inhibition. **A,** Western blot confirming expression of exogenous FLAG-tagged AXL in NCI-H358 cells. **B,** Comparison of $GI_{50}$ values for G12Ci, RASi, and adagrasib in NCI-H358 cells overexpressing AXL or GFP. The last column indicates the resistance ratio, calculated as $GI_{50}$ AXL / $GI_{50}$ GFP. **C,** IHC image of AXL expression in H358-GFP (top row) compared to H358-AXL (bottom row) **D,** Proliferation of NCI-H358 cells overexpressing GFP or FLAG-tagged AXL, measured as confluency in % of surface area over 96 hours. **E,** Body weight (in g) of mice treated with the indicated inhibitors over the course of the treatment period (corresponding to Fig. 4D). **F,** Statistical overview of tumor volume change ratios upon indicated inhibitor treatments (corresponding to Fig. 4D). Statistical significance was determined by two-way ANOVA followed by Tukey's multiple comparison test (n.s. = not significant, ***P* <0.01, *****P* < 0.0001).

**Supplementary Figure S5**

**A**

| HPAF-II CDX | | | |
|---|---|---|---|
| Median survival (days) | | | |
| Ctrl. | RASi | AXLi | Comb. |
| 12.5 | 16 | 15 | 34.5 |

**B**

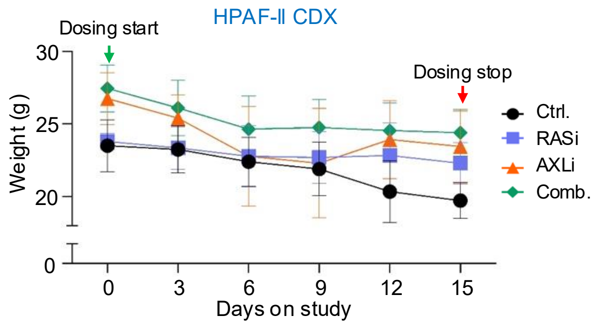


**C**

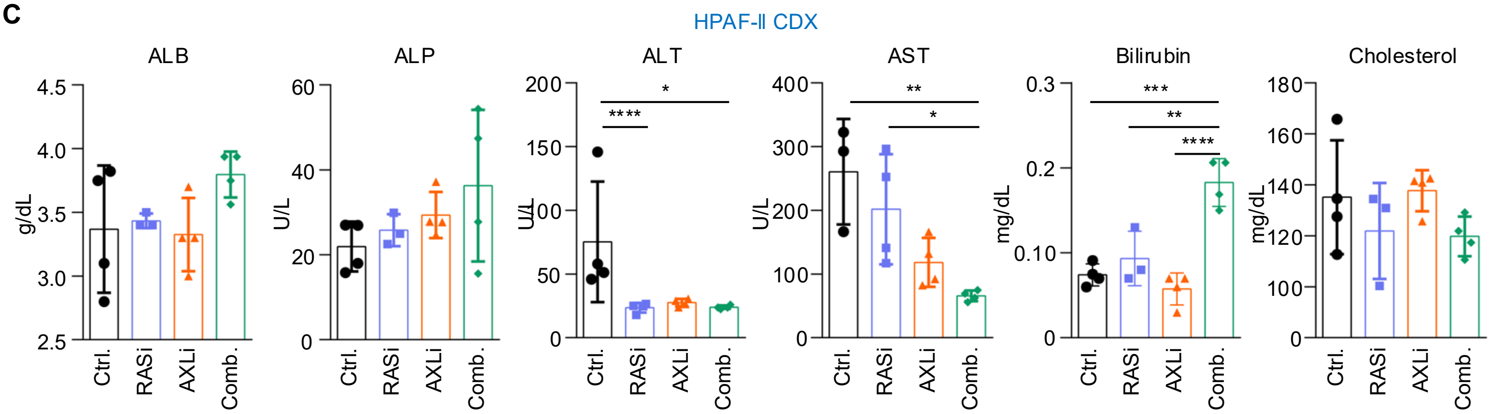


**D**

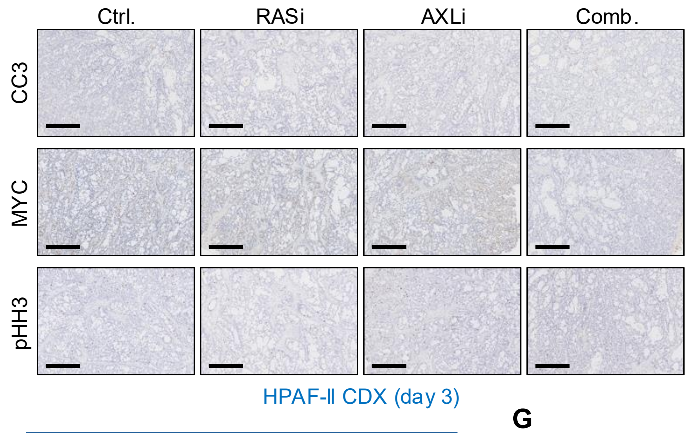


**E**

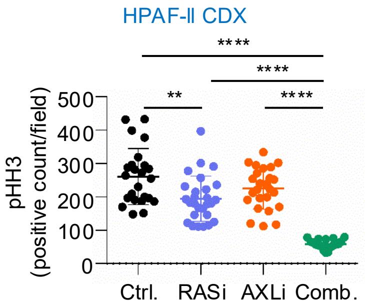


**F**

| KP-4 CDX | | | |
|---|---|---|---|
| Median survival (days) | | | |
| Ctrl. | RASi | AXLi | Comb. |
| 7 | 14 | 10 | 27 |

**G**

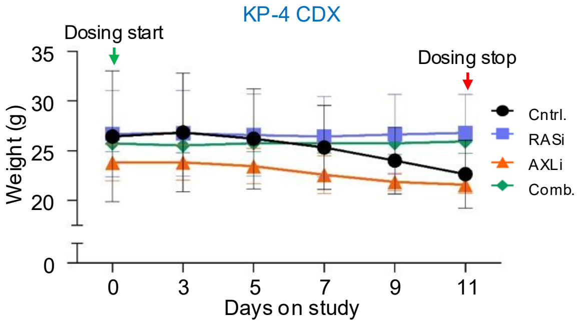


**H**

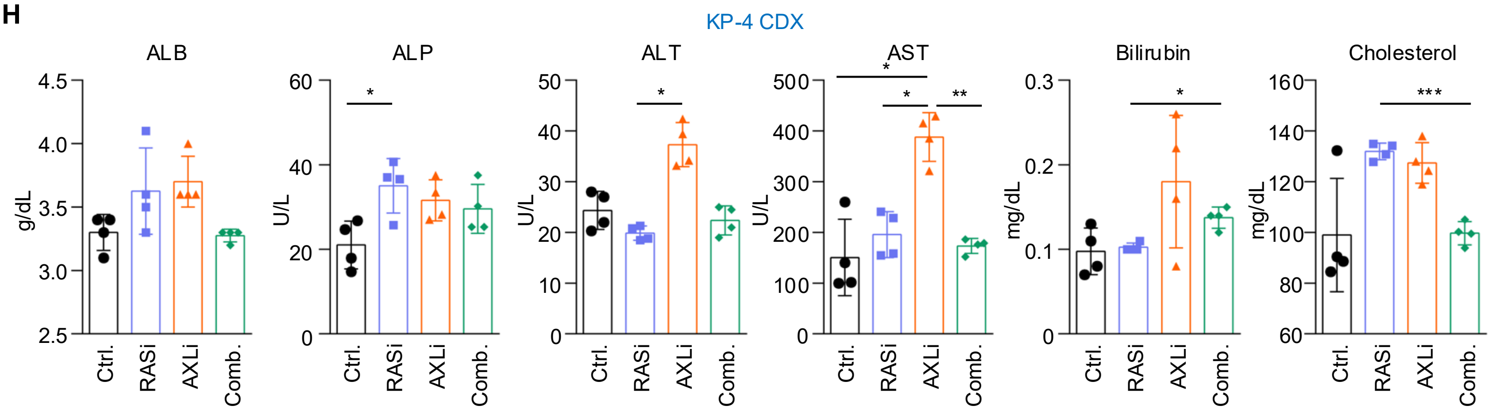

**Supplementary Figure S5**

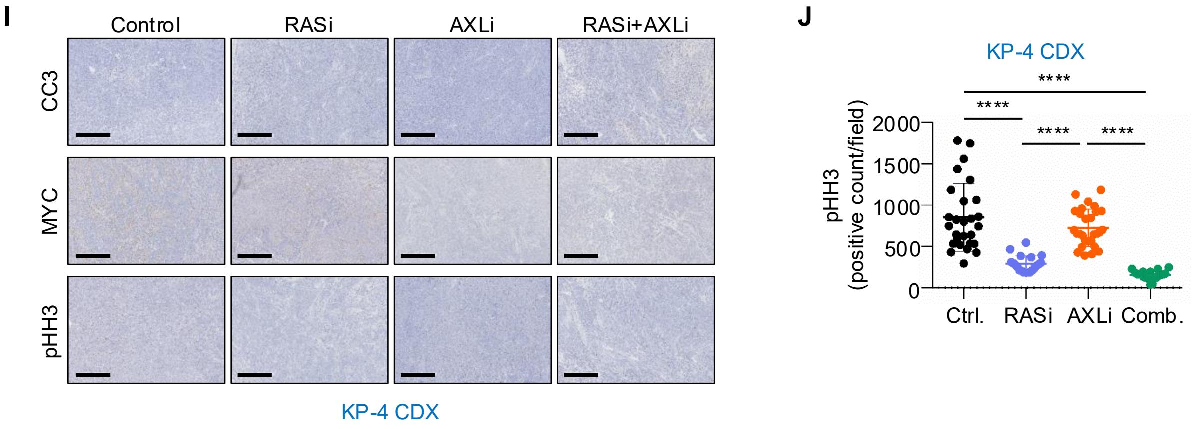


**Supplementary Figure 5.** AXLi enhances RASi antitumor efficacy in RASi-sensitive and -resistant PDAC CDX models. **A,** Table summarizing median survival in days of HPAF-II CDX tumor-bearing mice following treatment with vehicle control (Ctrl.), RASi, AXLi, or the combination (Comb.). **B,** Body weight changes of mice bearing HPAF-II CDX tumors treated with the indicated inhibitors over the course of the 15-day treatment period (corresponding to Fig. 5A) **C,** Liver toxicity assessment based on serum levels of albumin (ALB), alkaline phosphatase (ALP), alanine aminotransferase (ALT), aspartate aminotransferase (AST), bilirubin, and cholesterol in mice bearing HPAF-II tumors treated with vehicle, RASi, AXLi, or the combination. **D,** IHC images of HPAF-II tumor sections stained for cleaved caspase-3 (CC3), MYC, and phospho-histone H3 (pHH3). The scale bar indicates 600 µm. **E,** Quantification of pHH3-positive cells by IHC in HPAF-II tumor sections from mice treated with vehicle control, RASi, AXLi, or the combination. **F,** Table summarizing median survival in days of CDX mice bearing KP-4 tumors following treatment with control, RASi, AXLi, or the combination. **G,** Body weight changes of mice bearing KP-4 CDX tumors treated with the indicated inhibitors over the course of the 15-day treatment period (corresponding to Fig. 5F). **H,** Liver toxicity assessment based on serum levels of albumin (ALB), alkaline phosphatase (ALP), alanine aminotransferase (ALT), aspartate aminotransferase (AST), bilirubin, and cholesterol in mice bearing KP-4 tumors treated with vehicle, RASi, AXLi, or the combination. **I,** IHC images of KP-4 tumor sections stained for CC3, MYC, and phospho-histone H3 (pHH3). The scale bar indicates 600 µm. J, Quantification of pHH3-positive cells by IHC in KP-4 tumor sections from mice treated with vehicle, RASi, AXLi, or the combination. Data shown as mean ± SD. Statistical significance was determined by one-way ANOVA followed by Tukey's multiple comparison test, (*$P < 0.05$, **$P < 0.01$, ***$< 0.005$, ****$P < 0.001$).

**Supplementary Figure S6**

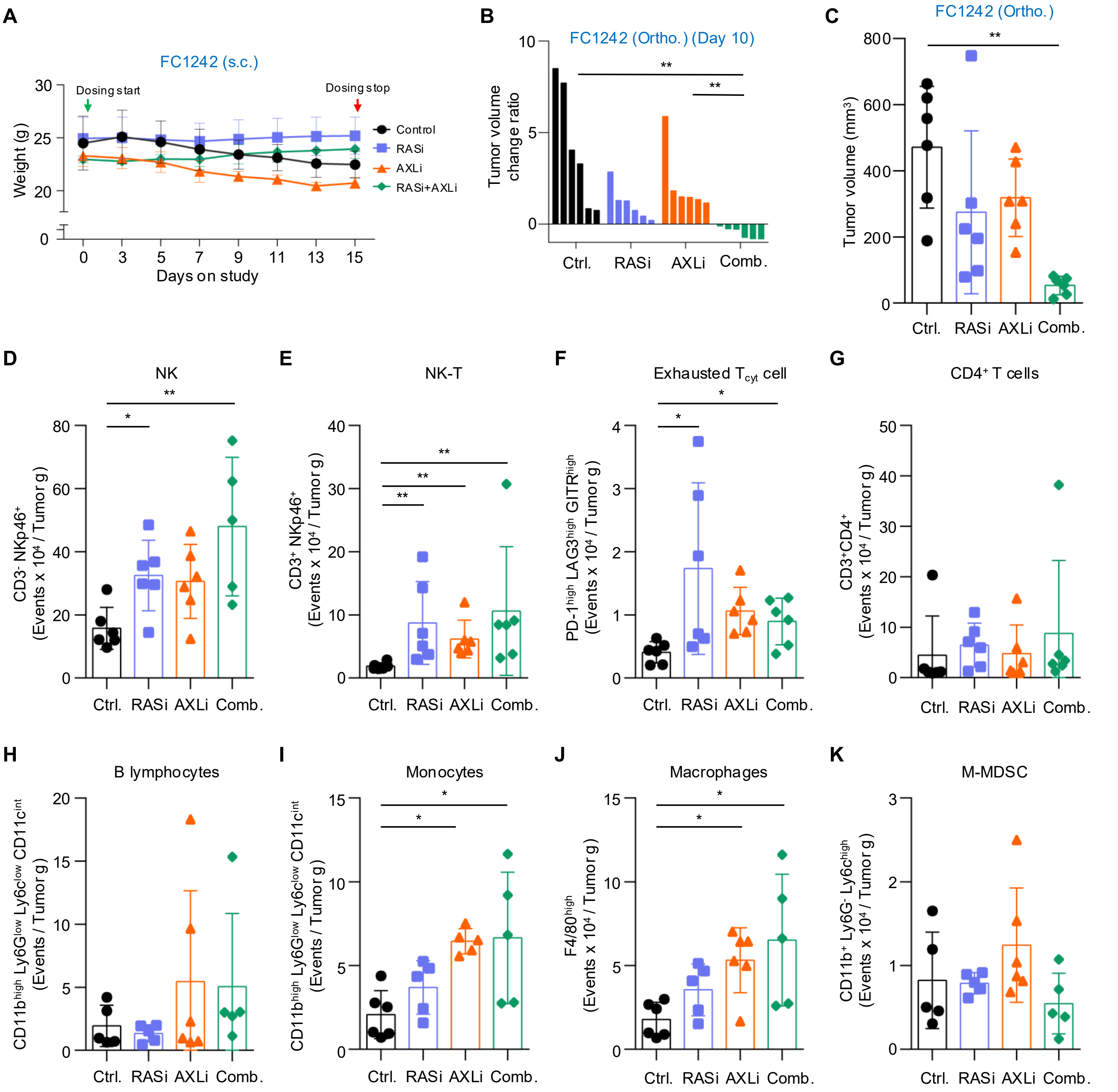


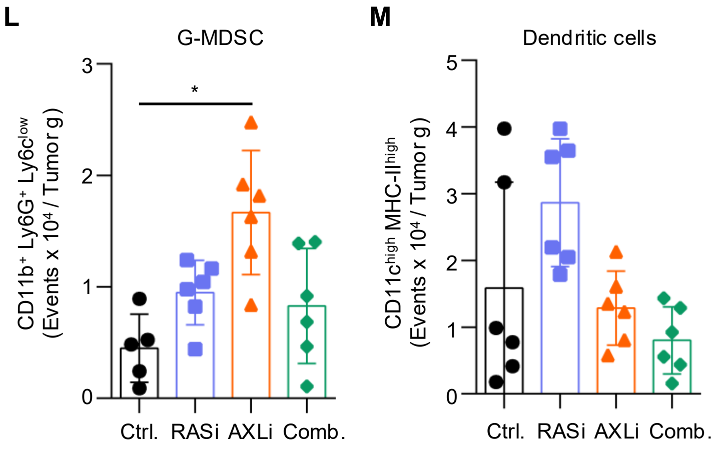


**N**

| Tumor volume change ratio | |
|---|---|
| **Comparison** | **Significance** |
| Ctrl. vs. AXLi | * |
| Ctrl. vs. RASi | **** |
| Ctrl. vs. Comb. | **** |
| AXLi vs. RASi | *** |

**Supplementary Figure 6.** Combined AXLi and RASi treatment extends survival and remodels the immune microenvironment in syngeneic PDAC models**. A,** Body weight (in g) of mice bearing subcutaneous KPC tumors treated with the indicated inhibitors over the course of the 15-day treatment period (corresponding to Fig. 6A). **B,** Waterfall plot showing the relative change in individual tumor volume from day 0 to day 10 (end of control group) for the FC1242 orthotopic model ($n$ = 6 tumors per group). Data shown as mean ± SD. Comparisons were made by Kruskal–Wallis test with FDR correction using the two-stage step-up method of Benjamini, Krieger and Yekutieli (*q < 0.05, **q < 0.01, ***q < 0.001, ****q < 0.0001). Only significant comparisons are shown.(**$P$ < 0.01). **C,** Absolute tumor volume ($mm^3$) for all four conditions; individual tumors of orthotopic FC1242 model collected at day 10 of treatment. Data shown as mean ± SD. Statistical significance was determined by one-way ANOVA followed by Tukey's multiple comparison test, (**$P$ <0.01). Only significant comparisons are shown. **D-M,** Immunoprofiling of tumor samples from C for **D,** NK cells **E,** NK-T cells **F,** Exhausted cytotoxic T cells **G,** $CD4^+$ T cells **H,** B lymphocytes **I,** Monocytes **J,** Macrophages **K,** M-MDSC **L,** G-MDSC **M,** Dendritic cells. Data shown as mean ± SD. Comparisons were made by Kruskal–Wallis test with FDR correction using the two-stage step-up method of Benjamini, Krieger and Yekutieli (*q < 0.05, **q < 0.01). Only significant comparisons are shown.**J**, Statistical overview of tumor volume change ratios upon indicated inhibitor treatments (corresponding to Fig. 6B). Statistical significance was determined by two-way ANOVA followed by Tukey's multiple comparison test, (*$P$ <0.05, ***$P$ < 0.001, ****$P$ < 0.0001).

### Supplementary Figure S7

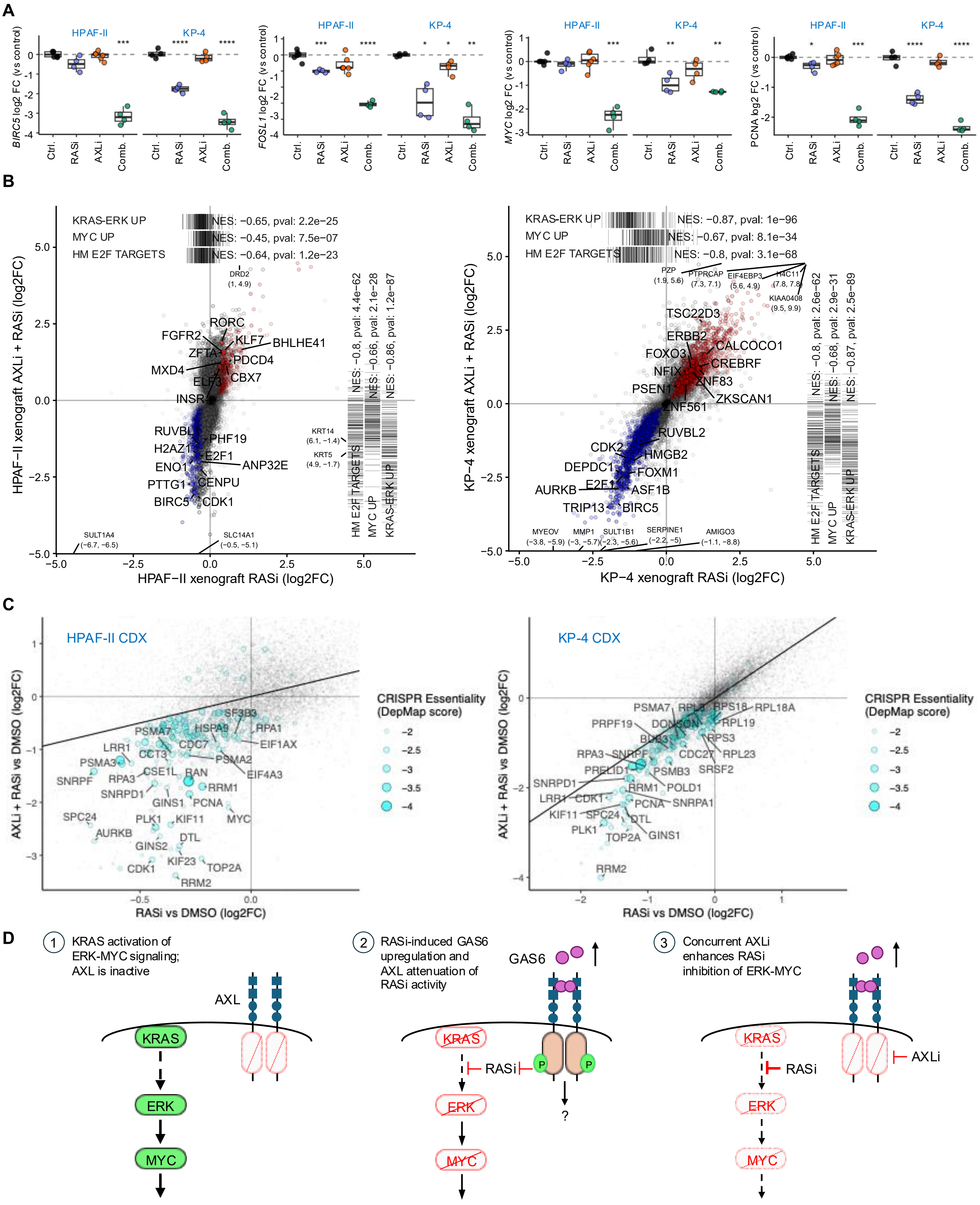

**Supplementary Figure 7.** Combination therapy with RASi and AXLi enhances RASi-dependent transcriptional changes in both RASi-sensitive and RASi-resistant CDX models. **A,** Box plots of RNA-seq from HPAF-II (left) and KP-4 (right) tumor samples, showing expression of *BIRC5* (Survivin), *FOSL1* (Fra1), *MYC*, and *PCNA* with indicated treatments. t-test *$P$ = 0., **$P$ = 0.01, ***$P$ = 0.001, ****$P \leq 0.0001$. **B,** Scatterplots showing RASi versus combination treatment in HPAF-II (left) and KP-4 (right) xenografts. Barcode plots on top and right indicate KRAS-ERK dependent gene sets, KRAS-ERK UP (1), MYC UP (2), and Hallmark (HM) E2F Targets (3). Highlighted genes are those significantly up (red) or down (blue) in both treatment groups. NES, normalized enrichment score. Labeled at periphery indicate genes falling outside of plot window. **C,** Differential dependency, RASi vs. combination, highlighting top essential genes from DepMap (CRISPR drop-out screen, smaller value = higher dependency). **D,** Proposed model showing the mechanistic basis behind the greater efficacy of the RASi and AXLi combination than of RASi alone: (1) Active RAS drives ERK-MYC signaling and AXL remains inactive; (2) loss of RAS-ERK-MYC signaling upon RASi treatment leads to feedback upregulation of GAS6 and activation of AXL, reducing inhibitory efficacy of RASi; (3) addition of AXLi enables the combination of RASi and AXLi to achieve deeper inhibition of RAS-ERK-MYC signaling.
